# The fate of social dilemmas under randomly fluctuating growth rates

**DOI:** 10.64898/2026.11.26.769452

**Authors:** Sanasar G. Babajanyan, Hrach G. Sargsyan, Armen S. Berujanyan, Kang Hao Cheong

## Abstract

We study three social dilemmas Prisoner’s Dilemma, Hawk-Dove and Coordination games between two phenotypes whose intrinsic growth rates switch randomly with the environment, modeled as dichotomous noise. We derive the exact stationary distributions of the population composition and the fixation probabilities. The persistence of each phenotype is set by its invasion rate, which depend only on the time-averaged environment, hence, zero-mean fluctuations do not alter the invasion abilities. Consequently, in the Prisoner’s Dilemma the takeover of defectors cannot be prevented unless the fluctuations of the intrinsic growth rates favor cooperators on average more than defectors. The switching rate of the environment, instead, shapes the distribution of the population composition, from bell-shaped to *M*-shaped and bimodal, and the fixation statistics of the Coordination game. Finally, since the growth rates and the payoffs enter the dynamics only in combination, different social dilemmas can be dynamically equivalent under different statistics of the fluctuations, so the underlying social dilemma can not be inferred from the population dynamics alone.

## INTRODUCTION

Evolutionary game theory provides a framework to study various phenomena across many disciplines, including but not limited to study natural selection, social behavior, and learning process [1–8]. The well-known and the simplest mechanistic description of the selection process between different phenotypes (strategies in game theoretic formulation) is provided by the replicator dynamics [5, 8, 9], whereby the more optimal phenotypes are passed through generations with greater success. The fitness of each phenotypes is assumed to be determined by pairwise interaction within and between different phenotypes.

The setup of the replicator dynamics assumes fixed environment for the individuals of given phenotype engaged in the pairwise interactions in the context of natural selection. Furthermore, it is assumed that the different phenotypes have the same intrinsic growth rates, such that the fitness difference between them is due to the payoffs. Equal intrinsic growth rates ensures the equivalence with the ecological formulation of the problem by means of Lotka-Volterra systems [5]. Relaxing the last assumption can change the evolutionary outcomes of the selection and differ from the predictions based on the payoffs [10, 11].

However, natural populations typically evolve in stochastic environments. The stochasticity of the environment may affect both the interactions and the intrinsic growth rates of individuals in phenotypically heterogeneous populations, such as tumors and microbial communities [12–16]. In a fluctuating environment, the evolutionary outcome can significantly differ from their counterparts in the fixed environment, both for periodic [17–23] and stochastic switching of the environment [24–35].

Here, we consider phenotypes whose intrinsic growth rates vary with fluctuating environment, either with equal means (symmetric fluctuations), or with one phenotype favored on average, through biased switching or unequal reproductive advantage. The environment randomly alternates between two states, modeled by dichotomous noise [36–38], favoring different phenotypes in each environment. The growth of the population size of each phenotype in the population is governed by the frequency-dependent fitness, which consists of the instantaneous growth rate and the payoff obtained from the game. The relative abundance of each phenotype is characterized by the modified replicator dynamics.

We consider the well-known 2 *×* 2 games, that is the social dilemmas, namely the Prisoner’s dilemma (PD), Hawk-Dove (HD) and Coordination (or Stag-hunt, SH) games with fluctuating instantaneous growth rates of different phenotypes engaged in the pairwise interaction. These simple games are of great importance for understanding of the evolution of social phenotypes at every level of biological organization [2, 39–58].

Fluctuations are known to promote cooperation through several mechanisms. In finite populations, environmental noise interacts with demographic noise and can strongly enhance the fixation probability of cooperators [26, 35, 59]. In discrete-generation model, payoff fluctuations compound multiplicatively, such that their variances and covariances enter the long-term growth rates, and cooperators can persist even under zero-mean fluctuations [60]. Similarly, in continuous-time models, stochasticity in payoffs or individual-level reproduction introduces variance-dependent terms into the replicator dynamics [29, 34].

Here, we consider a large population, so that demographic noise is negligible, in continuous time, with environ-mental fluctuations of finite magnitude and correlation time that are common for all individuals. The persistence of each phenotype is characterized by its invasion rate [61], and the long-term outcome through the stationary distri-bution of the population composition and the fixation probabilities. We show that in this setting the invasion rates depend only on the time-averaged environment, that is the first moment of the fluctuations, and independent of the characteristic timescale of them. Consequently, zero-mean fluctuations cannot prevent the takeover of defectors in the PD. Here, cooperators persist only if the environmental fluctuations effectively transform the PD settings to HD game.

The invasion rates, however, do not determine the shape of stationary distribution, which changes from bell-shaped to *M* -shaped and bimodal depending on the relative timescale of environmental switching and selection. Even if the invasion rates of phenotypes have the same as in case of averaged game, the fixation properties are altered by the stochasticity of the environment, for example in SH game. Finally, the noise in the intrinsic growth rates can produce the same long-term outcomes for different social dilemma, for example HD under symmetric fluctuations is dynamically equivalent to PD with unequal growth rate advantage. Hence the underlying dilemma can not be inferred based on the dynamics of the population composition alone in fluctuating environment.

## RESULTS

### Model: Replicator dynamics with fluctuating intrinsic growth rate difference

We consider frequency-dependent population growth model. We assume that the growth of each sub-population with fixed trait is given

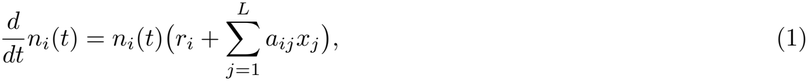

where *n*_*i*_(*t*) is the population size of phenotype *i* = 1, ..*L*, 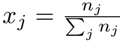 is the fraction of individuals of phenotype *j* = 1, ..*L* in the population, *a*_*ij*_ characterizes the symbiotic relationship between the phenotypes, that is the payoff matrix, and *r*_*i*_ is the intrinsic growth rate of phenotype *i*.

The time-evolution of the frequencies of each trait in the population, that is 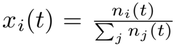 , is obtained from

(1) by using quotient rule, and is given by the following system of differential equations:

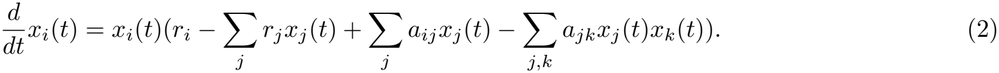

Note, that for equal reproduction rates, that is *r*_*i*_ = *r*_*j*_ for *i, j* = 1..*L*, we get replicator dynamics for time-evolution of the trait frequencies in the population.

For the considered case of two phenotypes in the population, defectors (*D*) and cooperators (*C*) for short, we obtain

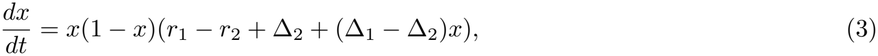

here, we used the normalization constraint on the frequencies and expressed the frequency of cooperators through that of defectors, *x*_2_ = 1 *− x*_1_ *≡* 1 *− x* (*x* is the fraction of defectors in the population). Δ_2_ = *a*_12_ *− a*_22_ and Δ_1_ = *a*_11_ *− a*_21_ are the payoff differences of the first and the second columns of the payoff matrix, respectively.

The stochasticity of the environment is incorporated in the model by assuming that the difference of the intrinsic reproduction rates of cooperators and defectors is time-dependent *r*_1_(*t*) *r*_2_(*t*) = *ϵξ*(*t*), where *ξ*(*t*) =*− ±* 1 is a symmetric dichotomous noise, *ϵ* shows the reproductive advantage and disadvantage of defectors over cooperators in favorable and unfavorable environment, respectively. Thus, the environment fluctuates between two states where opposite phenotypes have reproductive advantage.

Environmental switching occurs randomly with rates *λ*_*±*_ = *λ*(1*∓δ*), where *λ*_+_ is the switching rate from favorable to unfavorable environment, while *λ*_*−*_ governs the opposite switching. *δ∈* [*−* 1, 1] controls the bias of the environment towards these states.

Under the stochastically switching environment the time-evolution of the trait frequencies is governed by the following piecewise-deterministic Markov process

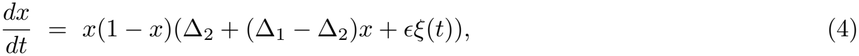

Equivalently to the above representation, one may assume that the reproduction rates are the same for both phenotypes, but each row of the payoff matrix is transformed to *a*_*ij*_ *a*_*ij*_ + *r*_*i*_ where *i, j* = 1, 2.

We study the behavior of the probability distribution function defined over the entire trajectory of the dynamics and over realizations of the fluctuating environment, that is *ρ*(*x*(*t*), *ξ*(*t*)). The behavior of *ρ*(*x*(*t*), *ξ*(*t*)) is given by the coupled forward Kolmogorov equations (see SI, section II.B for derivation), and in the stationary state the marginal distribution of the population composition alone is given by the following expression

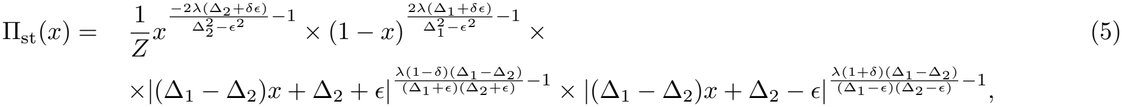

where *Z* is a normalization constant. In the main text we focus mostly on the unbiased environment case, that is *δ* = 0. In (5), each base corresponds to one of the equilibrium states of (4) in favorable and unfavorable environment, while the exponents control the relative likelihood of observing the given equilibrium state in the stationary trait distribution. The stationary distribution is found by considering the games that exhibit one stable equilibrium on the domain of all possible *x ∈* [0, 1] in each environment.

However, for the stag-hunt games this approach is not directly applicable since the coexistence of both phenotypes is not evolutionarily stable in each environment. We discuss this issue in detail below and in SI, section III.B.

We also discuss the invasion rates of each trait when rare showing whether small population of a given trait will grow in a resident population composed of only by its competitors. The invasion rate of *D* trait in a resident population composed of only *C* for stochastically switching environment is given by the following expression [61]

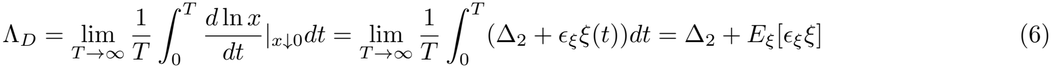

similarly, the invasion rate of *C* in a resident population composed of only *D* is

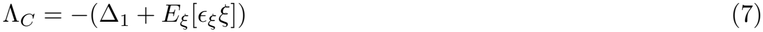

The positive invasion rate means that a trait can invade the resident population when rare. Equations (6) and (7) are written for the general case, that is the reproductive advantage and disadvantage of *D* trait in favorable and unfavorable environment are not equal in magnitude *ϵ*_+1_ ≠ *ϵ*_*−*1_. The average is taken over the stationary probabilities of environmental fluctuations, which need not be dichotomous in general. The limit w.r.t. time of the environmental fluctuations is replaced by its mean due to the ergodicity of the fluctuations.

For *ϵ*_+1_ = *ϵ*_*−*1_, but biased environmental fluctuations, we get that the invasion rate of *C* and *D* phenotypes are equal to Λ_*C*_ = *−*(Δ_1_ + *ϵδ*) and Λ_*D*_ = Δ_2_ + *ϵδ*, respectively, since *E*[*ξ*] = *δ*. Note, that the invasion rates Λ_*C*_ and Λ_*D*_ are involved in the exponents of the terms with base *x* and 1*− x* in Π_st_(*x*), given by (5). The long-term evolutionary outcomes, though, are not fully characterized by the invasion rates, that is various distributions of the population composition can be observed for the given sign of invasion rates.

For the HD and Stag-hunt games we consider unbiased environmental fluctuations, *δ* = 0, and equal intrinsic growth rate advantages of each trait in their favorable environments, *ϵ*_+1_ = *ϵ*_*−*1_. The other cases are discussed within the context of PD game.

In the HD game (see Fig.1A), the payoff differences are such that Δ_1_ *<* 0 *<* Δ_2_ resulting into an evolutionarily stable state where both phenotypes coexist with frequencies 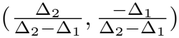 and the replicator dynamics converges to that stable equilibrium state starting from initially heterogeneous population, *x*_0_ *∈*(0, 1). Thus, both cooperators and defectors can invade the population when rare in the absence of environmental fluctuations.

**FIG. 1:**
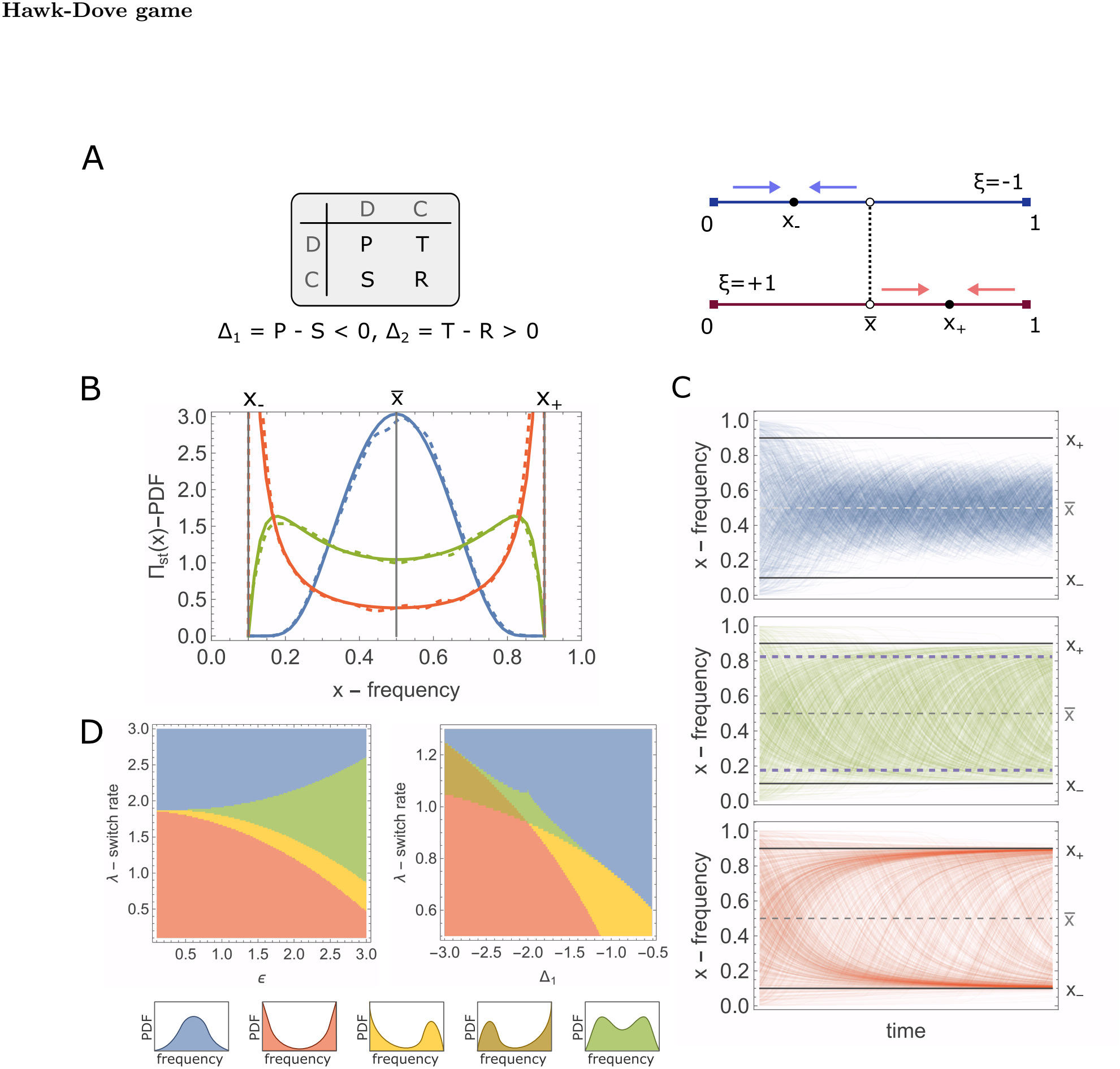
Evolutionary outcomes for the Hawk-Dove game with fluctuating intrinsic growth rates for different model parameters. **A)**The payoff matrix structure and sign of payoff differences in HD games, right panel. evolutionarily stable states for unfavorable (*x*_*−*_), favorable (*x*_+_) and averaged environment (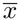, obtained for *ϵ* = 0 in (4)). Blue and red arrows show direction of the selection process in the unfavorable and favorable environments, respectively. **B)**Solid and dashed lines show the distribution of the population composition at stationary state obtained theoretically, (5), and via simulations of (4) (see details in Method section), respectively. Blue, Green, and Red curves corresponds to fast, intermediate (*M* -shaped distribution) and slow switching regimes of the environment, respectively. **C)**From top to bottom, trajectories of *x*(*t*) for fast, intermediate and slow environment-switching regimes, whereby the typical environmental episode lasts 0.2, 1.25 and 5 times the population relaxation time towards the ESS of the averaged game, respectively. The model parameters for B) and C) are Δ_1_ = *−*1.0, Δ_2_ = 1.0, *ϵ* = 0.8, *λ*_1_ = 2.5 (blue), *λ*_2_ = 0.4 (green), *λ*_3_ = 0.1 (red). **D)**Phase-space of the possible stationary distribution for various model parameters. The color of each region shows the shape of the stationary distribution presented on panels below. The remaining model parameters for left and right panels are Δ_1_ = *−*4.0, Δ_2_ = 3.5 and Δ_2_ = 2.0, *ϵ* = 0.5, respectively.

In the fluctuating environment, the fraction of defectors at the equilibrium state, in both favorable and unfavorable environments, is shifted due to the advantage and disadvantage in the intrinsic reproduction rates, respectively. The fluctuations in the intrinsic growth rates do not alter the invasion rates of both phenotypes when rare, (6,7), since *E*_*ξ*_[*ξ*] = 0.

The stable equilibrium states are 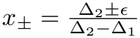 in a favorable and unfavorable environment, respectively, where the intrinsic growth rate difference must satisfy *ϵ <* min[|Δ_1_|, Δ_2_] to ensure *x*_*±*_ *∈* [0, 1]. Due to the unique evolutionarily stable states in each fixed environment, *ξ* = *±*1, the support of the trait distribution at stationary state, (5), is the interval bounded by these states, [*x*_*−*_, *x*_+_].

Nevertheless, it is still possible that neither of evolutionarily stable states in a fixed environment, that is *x*_*±*_, will likely be observed as an evolutionary outcome. We consider the case of equal payoff difference by modulo, that is, Δ *≡ −*Δ_1_ = Δ_2_ for sake of simplicity. In this case, the evolutionarily stable states are 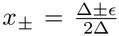 . Thus, in a favorable and unfavorable environment, the fraction of *D* in the evolutionarily stable increases and decreases by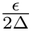, respectively.

Analyzing the stationary probability of the population composition, (5), reveals possible evolutionary outcomes. In particular, considering the exponents of Δ(1 *−* 2*x*) *± ϵ* terms in (5), it follows that neither of the evolutionarily stable states is likely be observed, Π_st_(*x*_*±*_) *→* 0 when environmental fluctuations are fast compared to selection,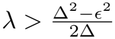. For 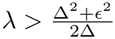, the distribution of population composition develops a unique peak at 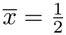(see Fig.1B and C for distribution and time-dependence of *x*), corresponding to ESS of the averaged environment (equivalent to *ϵ* = 0 in (4)). Increasing *λ* narrows the distribution of *x* around 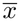.

The fast-fluctuating environmental regime does not necessarily produce unimodal distributions around 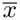. Indeed, 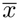can also be a local minimum, whenever 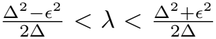 . In this regime, the trait distribution develops two symmetric peaks around 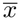, resulting in a *M* -shaped distribution (see Fig.1B,C and SI, section III.A for further details on derivation). Thus, it is unlikely to observe ESS corresponding to fixed or averaged environments *x*_*±*_ and 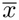, respectively (see Fig.1B).

In contrast to fast-fluctuating regime, the evolutionarily stable states *x*_*±*_ become the most likely outcomes when the environmental fluctuations are slow compared to the selection process,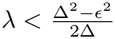, whereby Π_st_(*x±*) *→ ∞* (but still integrable). Thus, selection under rare fluctuations of the environment produce bimodal distribution of the population composition (see Fig.1B,C).

So far we assumed equal in magnitude payoff differences, that is, |Δ_1_| = Δ_2_ = Δ. Therefore, if |Δ_1_| *≠* Δ_2_ it is possible to get different likelihoods of observing evolutionarily stable states of the fixed environment *x*_*±*_, which is impossible in a symmetric case. Indeed, for Δ_1_ *<* 0, Δ_2_ *>* 0 and 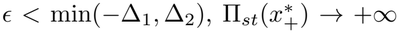 and the integral converges near *x*^*∗*^ whenever 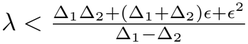. Similarly, 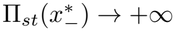 and the integral converges near 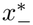 if 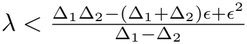 (see SI, section III.A). The possible evolutionary outcomes for asymmetric payoff differences is shown in Fig.1D. Increasing *λ* for fixed *ϵ* results in noise-induced transitions in the trait distribution, from bimodal to *M* shaped distributions, and consequently into unimodal distributions. However, in contrast to the case of equal payoff difference, here it is possible to observe a local maxima in the trait distribution which is not the most likely outcome, since the trait distribution diverges on the opposite endpoint of the local maxima (yellow and brown regions in Fig.1D).

### Dominance Game: Prisoner’s Dilemma

In the PD game, the payoff differences have the same sign, that is Δ_1_ = *P− S >* 0 and Δ_2_ = *T− R >* 0, resulting into takeover of defectors in averaged game, which is the only ESS in that case (see SI, section III.C for the short review).

The invasion rate of *C* is negative in symmetric fluctuating environment, Λ_*C*_ *<* 0, hence, the cooperators can not invade the resident population of all *D*. Furthermore, in the long-run they extinct, even if initially present with non-negligible fraction.

Indeed, in the vicinity of *x* = 1, the stationary distribution behaves as 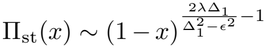 , here we consider Δ_1_ *< ϵ <* Δ_2_, but the results hold for all other orderings as well (see SI section III.B for case studies). The exponent in the last expression is always smaller than *−*1, hence, the divergence is non-integrable. Meanwhile, close to fluctuation-induced equilibrium state 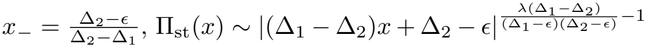, which is integrable since for all *λ* the exponent is always greater than *−*1. The symmetric fluctuations of the environment does not prevent the takeover of *D* in the population, since they suffer less in unfavorable environment than *C* does due to the payoff advantage over *C* for all possible composition in the population.

Although the takeover of the defectors is inevitable under symmetric environmental fluctuations, the mean time necessarily for the takeover behaves differently for various payoff differences. For the given initial state and environmental switching rate, increasing Δ_1_ increases the mean extinction time of cooperators, whereby the fraction of cooperators drops to threshold value close to zero, compared to the deterministic case with no difference in the reproduction rates of *C* and *D*. For smaller Δ_1_ the stochasticity of the environment accelerates the extinction of *C*, while no such a dependence observed for Δ_2_ (see SI, section III.B for details).

Thus, for the survival of cooperators the environment must favor them more than defectors on average. That is, it is necessary for cooperators to be able to invade the population of defectors when rare, Λ_*C*_ *>* 0.

In what follows, we discuss two different scenarios that favor *C* more than *D*, such that the takeover of *D* can be prevented.

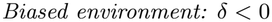

The invasion of cooperators is possible in the biased environment with equal in magnitude intrinsic growth rate difference in favorable and unfavorable environment, respectively, whenever the environment is biased towards *C*-favored state, *δ <* 0, such that

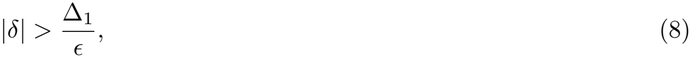

The condition (8) follows from Λ_*C*_ = *−*Δ_1_ *− ϵδ >* 0, thus, the cooperators can invade the population of all *D*. The invasion of *C* is independent of Δ_2_ = *T − R* and *λ*. For the survival of *D*, on the other hand, it is necessary that Δ_2_ *> ϵ*|*δ*|, which always holds for the considered ordering of the model parameters Δ_1_ *< ϵ <* Δ_2_.

Invasion of *C* is equivalent of considering an averaged game with payoff difference 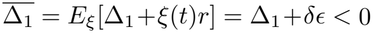, corresponding to HD game. Selection drives the fraction of *D* towards coexistence induced by intrinsic growth rate difference, *x*(*t*) *→ x*_*−*_, and the takeover state, *x* = 1, in unfavorable and favorable environments, respectively, the former played more than later on average.

We observe different evolutionary outcomes in these biased environment satisfying (8) (see Fig.2B left panel). Namely the bell-shape distribution is observed when 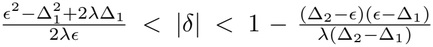 (see Fig.2B right panel yellow curve for distribution function and Fig.2C for trajectories). For |*δ*| below and above the interval the probability distribution peaks on the endpoints of the support, that is, only *D* and *x*_*−*_, respectively (see Fig.2B right panel and Fig.2C green and red curves, respectively).

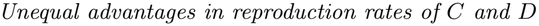

**FIG. 2:**
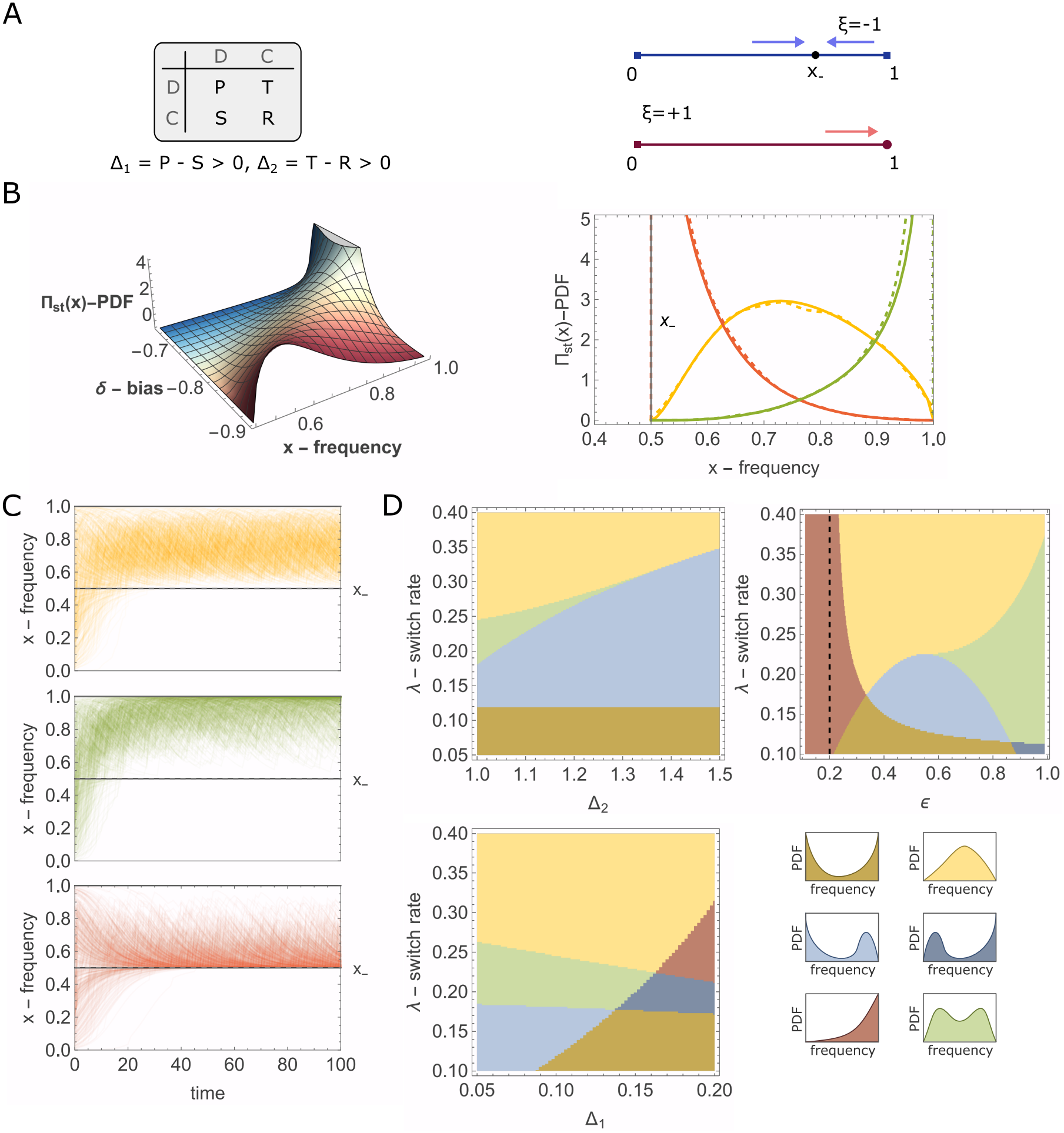
Evolutionary outcomes for Prisoner’s Dilemma game in fluctuating environments that are either biased or provide unequal advantages in intrinsic growth rates. **A)**Payoff-matrix of the game and ESS in an unfavorable and favorable environments as in Fig.1A. **B**) Probability distribution of the population composition for various model parameter in biased environments favoring *C*. The left panel shows the probability distribution for different values of the environmental bias. In the right, yellow curve (*δ* = *−*0.83) represents a scenario where the likelihood of observing stable equilibrium states of the fixed environments at stationary nullifies. Red (*δ* = *−*0.95) curve shows the case where the coexistence of *C* and *D* with fractions (*x*_*−*_, 1*− x*_*−*_) is the most likely outcome. Green curve (*δ* =*−*0.7), in contrast, the stationary distribution is concentrated near the all-*D* boundary, although cooperators persists. The remaining model parameters are Δ_1_ = 0.5, Δ_2_ = 1.0, *ϵ* = 0.75, *λ* = 2.0. **C)**Trajectories of *x*(*t*) obtained from the simulations for the model parameters of Fig.2B right panel, with respective colors. **D)**Phase-space of the probability distribution of the population composition at the stationary state for unequal advantages in intrinsic growth rates of cooperators and defectors. The colors correspond to regions of probability distributions of different shapes, these are presented in the small panels on bottom-left. The dashed black line corresponds to *ϵ* = 2Δ_1_ value, (9), that separates the regions corresponding to extinction of cooperators and their survival in small fractions, though. The remaining model parameters are Δ_1_ = 0.1, *ϵ* = 0.75 (top left), Δ_1_ = 0.1, Δ_2_ = 1.0 (top right), and Δ_2_ = 1.0, *ϵ* = 0.75 (bottom left).

Another scenario that prevents takeover of cheaters is the unbiased environmental fluctuations, *δ* = 0, with greater intrinsic growth rate advantage of *C* over *D* in the environment favoring cooperators. For sake of simplicity, we assume that *D* has no intrinsic growth rate advantage over *C* in *ξ*(*t*) = 1 environment,*ϵ*_+_ = 0, while in *ξ*(*t*) = *−*1 environment *C* has growth rate advantage of *ϵ*_*−*_ = *ϵ*. The invasion rate of cooperators, Λ_*C*_, is positive whenever

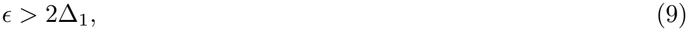

and the stationary distribution of the population composition has the following form (see SI, section III.B for derivation)

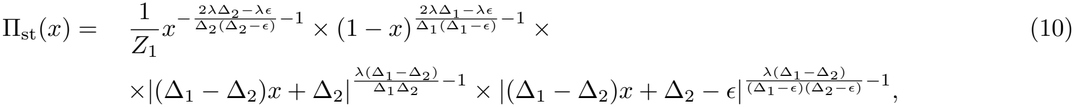

The distribution (10) can only diverge (non-integrable divergence) close to *x* = 1 point, that is the takeover of cheaters, which is prevented by the condition (9).

The case of unequal advantage in intrinsic growth rates is equivalent to that of symmetric noise (unbiased and equal advantage), where the payoff differences are 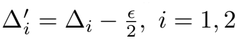 and the intrinsic growth rate advantage is 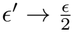 (see SI, section III.B for derivation). Then, the condition (9) is equivalent to 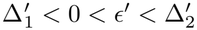, which is an ordering of HD games, whereby the stable ESS are *x* = *x*_*−*_ and *x* = 1 in unfavorable and favorable environments. Thus, the unequal advantage in PD game can be analyzed by the means of HD game. As in the HD game, here, the evolutionary outcomes span all possible regimes, from bell-shape distribution in fast-switching regime, to the dominance of one trait and *M* shaped distributions (in intermediate regimes of environmental fluctuations), to bimodal distribution in the slow-switching regime of the environment.

Thus, in both cases discussed above, the takeover of defectors is prevented if the averaged game corresponds to that of HD game, for which Λ_*C*_ = *−*(Δ_1_ + *E*_*ξ*_[*ϵ*_*ξ*_*ξ*]) *>* 0. In more general case of environmental fluctuations, the takeover of defectors in the population is prevented for

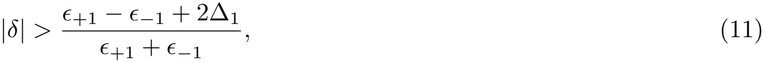

such that *δ <* 0, independent of Δ_2_ and *λ*.

### Coordination (Stag-Hunt) game

In SH games, the sign of payoff differences is the opposite of that in HD games, that is, Δ_1_ *>* 0 and Δ_2_ *<* 0. Here, with equal intrinsic growth rates *r*_1_ = *r*_2_, neither trait can invade when rare, and the evolutionarily stable states are corresponding to homogeneous population composed of only *D* or only *C* traits. These equilibrium states are separated by the unstable equilibrium 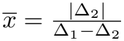 of the replicator dynamics (3). Thus, for 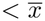 and 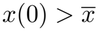 the population evolves toward only *C* and only *D* states, respectively.

In general, the SH game can be transformed to effectively PD game, that is the invasion rates have different signs, but the mutual invasion is impossible, that is Λ_*C*_, Λ_*D*_ *>* 0 impossible.

We focus here on the symmetric fluctuating-environment, since it resembles the selection process in SH game in the absence of intrinsic growth rate difference, but alters the fixation statistics of the phenotypes.

The invasion rates Λ_*D*_ and Λ_*C*_ are negative for both traits, hence, invasion through symmetric environmental fluctuations is impossible. The homogeneous states of the population, corresponding to only *D* and only *C* states, remain stable in both favorable and unfavorable environments, as follows from the stability analysis of (4) for *ξ* = *±*1 (see SI, section III.C for the derivation).

Therefore, the difference in the intrinsic growth rates alters the position of the unique unstable equilibrium state in each environment, thus altering the fixation statistics of each trait. In an unfavorable and favorable environment the unstable equilibrium state shifts to 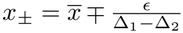, respectively. These equilibrium states are feasible, that is 0 *< x±<* 1 , for 0 *< ϵ <* min[Δ_1_ , *−*Δ_2_].

For any trait composition in the population below *x*_+_, *x*(*t*) *< x*_+_, and above *x*_*−*_, *x*(*t*) *> x*_*−*_, the fixation of *C* and *D* traits are inevitable, respectively, the selection gradients in each environment point at the same direction in these regions (see Fig.3A right sub-panel).

**FIG. 3:**
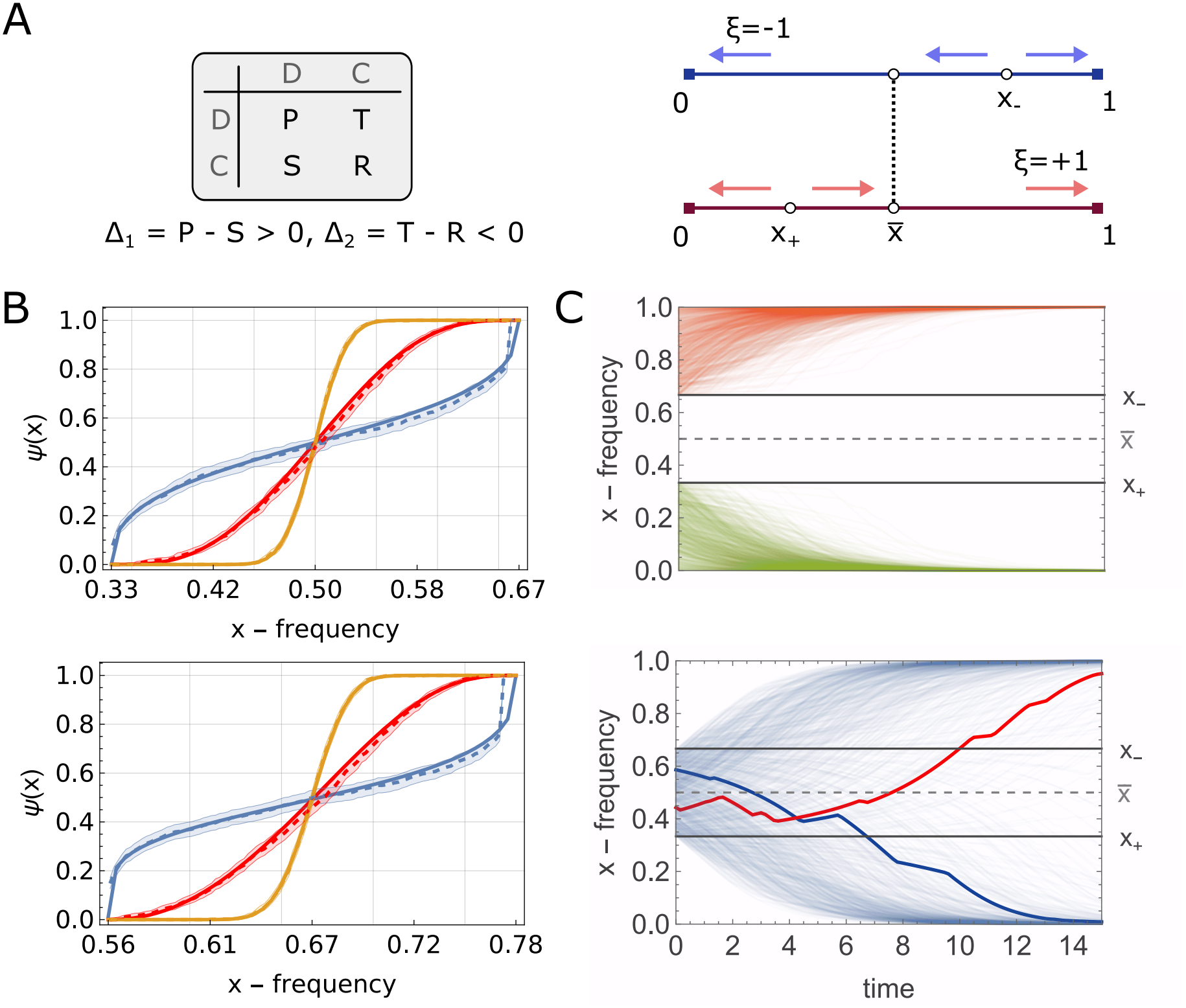
Evolutionary outcomes for the Coordination, or Stag-Hunt game, for various environmental switching regimes. **A)**Left and right panels show payoff matrix structure and ESS for favorable, unfavorable and averaged environment, analogous to that of Fig.1A. **B)**Fixation probabilities of trait *D* starting from the region (*x*_+_, *x*_*−*_), where the selection has opposite directions in favorable and unfavorable environments. Blue, red and orange curves show the fixation probabilities obtained theoretically (full) and from simulations (dashed) lines for slow (*λ* = 0.1), intermediate (*λ* = 1) and fast (*λ* = 10) environmental switching regimes, respectively. The shaded areas show the two-sided 95% Wilson score confidence intervals around the sample mean for the given values of the model parameter. In the top and bottom panel,Δ_1_ = *−*Δ_2_ = 0.6, *ϵ* = 0.2 and Δ_1_ = 0.6, Δ_2_ = *−*1.2, *ϵ* = 0.2, such that 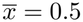 and 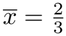 , respectively. **C)**Trajectories of *x*(*t*) for various initial states from outside (top) and inside (bottom) the region (*x*_+_, *x*_*−*_). Blue and red trajectories show the fixation of a trait with initial states favoring opposite traits. The model parameters are the same as red curve of top panel in B).

The non-trivial scenario in the evolutionary outcomes occurs for the population composition in between the unstable equilibrium states, that is, *x*_+_ *< x*(*t*) *< x*_*−*_, which is only possible if initially the population composition is from the same interval *x*(0) *∈* (*x*_+_, *x*_*−*_).

To determine the evolutionary outcomes for the population composition in the interval (*x*_+_, *x*_*−*_) we introduce two artificial absorbing boundaries located at *x*_+_ + *l* and *x*_*−*_ *− l*, respectively, where *l ∼ o*(1) [62–64], such that switching of the environment is unlikely in the interval *l*. Let *Ψ*_*ξ*_(*x*) denote the probability that, starting with the initial composition *x* (*x∈* [*x*_+_ + *l, x*_*−*_ *l*]) in the environment *ξ* = *±*1, the population composition reaches the right boundary before reaching the left one, hence, enters the region of inevitable fixation of *D*. For an unbiased environment, the mean probability of reaching the right boundary irrespective of the initial state of the environment is given by 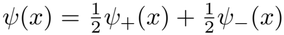 Note, that crossing the given unstable equilibrium *x*_*±*_, hence, ending up in either *C* or *D* states, is possible only if the environment is in opposite state.

The probability *Ψ*(*x*) is presented in Fig.3B for various payoff differences, Δ_1_ and Δ_2_, and for different environmental switching rates *λ*.

In the unstable equilibrium of the SH game in the averaged environment, 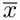, the fixation of *C* or *D* are equally likely, that is 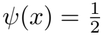 , for Δ_1_ = *−*Δ_2_, irrespective of *λ* (see Fig.3B). Indeed, the position of 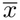 depends on the payoff differences, however, the boundaries are located at equal distances from 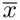 , and given the unbiased environment, *δ* = 0, it is equally likely to reach either of the boundaries started from 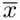.

Increasing *λ* sharpens the curve *Ψ*(*x*), and in the limiting case *λ→ ∞* approaches to the deterministic of fixation properties in averaged environment, that is, *Ψ*(*x*) = 0 and *Ψ*(*x*) = 1 for 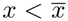 and 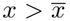, respectively. Indeed, for larger *λ* the stochasticity of the environment self-averages to zero, since it is unbiased, and the selection process behaves deterministically. Decreasing *λ*, smoothens the curve of fixation probabilities, whereby a trait currently in minority can fixate in the population with greater chance. Indeed, smaller *λ* implies greater persistence time of the current environment, hence, the population can evolve to either boundary in the current environment. Thus, for *λ→* 0, the fixation of either trait will resemble the statistics of environmental fluctuations, for unbiased environment the fixation of either trait become equally likely.

The trajectories of independent runs for intermediate switching regime are illustrated in Fig.3C. The trajectories starting outside of (*x*_+_, *x*_*−*_) approach the closest phenotypically homogeneous equilibrium state of the population. Therefore, the trajectories starting inside the interval (*x*_+_, *x*_*−*_) do not necessarily reach the initially nearest equilibrium state (blue and red curves), in contrast to deterministic case with 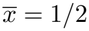.

## DISCUSSION

We study the evolutionary outcomes of social dilemmas in the population consisting of two phenotypes, namely, cooperators, *C*, and defectors, *D*, in a stochastic environment that alternates between two states favoring the reproduction of one phenotype over the other, while neglecting the demographic noise in the model by assuming large population. We assume that the growth of the population of each trait is governed by the frequency-dependent fitness consisting of terms of pure reproduction and competition for common resources, that is represented by the payoff matrix, Hawk-Dove, Prisoner’s Dilemma or Coordination game.

The fluctuations in the intrinsic growth rates can change drastically the long-term outcomes of the selection process. We analyze the evolutionary outcomes of each social dilemma by means of average invasion rate of each trait and stationary distribution of the population composition.

The invasion rates define whether in the long-run the given trait will persist in the resident population composed of only competitors. Environmental fluctuations can induce invasion of a trait that, otherwise, has no chance to invade in the fixed environment. Thus, the social dilemma can be transformed to effectively another one, such as from PD to HD or SH to PD games. Due to these transitions the coexistence or takeover of a trait that is not favored by the selection in the absence of environmental fluctuations is possible.

The invasion rates, however, do not predict the shape of the distribution of population composition at stationary state. To this end, the coupled Kolmogorov equations are solved for HD and PD games.

In HD game, for fast-fluctuating symmetric environment we observe bell-shape distribution around the equilibrium of HD game with no growth rate difference between *C* and *D* phenotypes. The probability of observing either evolutionarily stable states, that are the endpoints of distribution function, of the selection in favorable and unfavorable environment, respectively, tends to zero. Increasing fluctuation rates further results in narrower distribution, thus resembling the deterministic selection process. For intermediate switching rates, an *M* shaped distribution develops, whereby neither ESS corresponding to fixed or averaged (no difference in intrinsic growth rates) is likely to be observed. Instead, noise-induced peaks are symmetrically developed around the ESS of averaged game. In the slow-environment, the ESS of fixed environments become the most likely outcomes to observe in the long-run. For asymmetric payoff differences, the distribution of the population composition can develop a heavy tail on one of the endpoints.

For the PD game, symmetric fluctuations of the environment do not prevent the takeover of the defectors, as follows from the invasion rate analysis of cooperators. Instead, the environmental fluctuations need to favor *C* either through its statistics of switching or through the unequal magnitudes in intrinsic growth rates, or both of them. We derive the general condition for preventing the extinction of cooperators in a fluctuating environment. The survival of cooperators is due to the transformation of PD to HD game. In the discrete generation model, the invasion rate corresponds to the geometric mean of the per-generation fitness ratio, which expansion contains covariance terms that can prevent the takeover of defectors in a symmetric fluctuating environment with zero-mean [60]. In the considered case, the per-capita growth rate depends linearly on payoffs, and the invasion rate is only dependent on the mean of environmental fluctuations. Hence, zero-mean fluctuations do not prevent the takeover of the defectors. The survival of cooperators, that is, fixation in finite population, is possible if the demographic noise is incorporated in the model, whereby the fluctuations of the total population size enhance the fixation of cooperators in the population [26, 35]. As in the HD, the distribution of the population composition spans from bell-shaped to *M* shaped and bimodal ones depending on the environmental fluctuations.

In case of Coordination game, we focus on the fixation statistics of the phenotypes, analyzed through coupled backward-Kolmogorov equations, in symmetric fluctuating-environment, whereby in each environment the invasion of any phenotypes is impossible, otherwise SH transforms to PD game. The phase-space structure of the game is preserved in both favorable and unfavorable environments, that is, all *C* and all *D* states are ESS in both environments which are separated by an unstable equilibrium state, whose position depends on the current environment. In between these environment-induced unstable equilibrium states the selection gradient alternates with switching environment, in contrast to other regions in the phase space where selection gradient has the same direction in both favorable and unfavorable environments. The fixation of either trait started from the region of opposite selection gradients is equally likely for the initial state corresponding to unstable equilibrium of the averaged game with equal payoff differences, which is always located in the middle of the region of opposite selection gradients. Equal chances of fixation of each trait holds in rarely switching unbiased environment as well. The fixation probabilities in this setup resemble that of averaged game by increasing the environmental fluctuation rate, which in the limiting case of self-averaged environmental noise leads to step-function behavior of fixation whereby the outcome is determined by the location of the initial state relative to an unstable equilibrium of the averaged environment.

In conclusion, we show that the fluctuations in the intrinsic growth rates may lead to rich phase-space structure of the evolutionary outcomes, even in the simplest case of a population with two traits. These fluctuations can induce transitions between qualitatively distinct evolutionary outcomes of social dilemmas and may even change the structure of the effective (averaged) game, through sign flipping of the invasion rates of the traits. The findings here are twofold. First, it is shown that by altering intrinsic growth rates the traits that are disadvantageous in a given social dilemma can outcompete or coexist with their competitors, these are important for antibiotic resistance and tumor proliferation, where evolutionary dynamics can inform therapeutic strategies. Second, the long-term outcome alone can not identify the underlying social interactions. Because only the environment-specific combinations of payoffs and intrinsic growth rate differences enter the dynamics; the PD game with unequal advantage in intrinsic growth rates is dynamically identical to a HD game in symmetric fluctuating environment. Separating the two requires independent measurements of the population growth in favorable and unfavorable environments, whereby comparing the log abundance ratio against phenotype frequency in each fixed environment and in fluctuating environment can reveal the underlying social-dilemma. Moreover, parallel curves of the log abundance ratio against the phenotype frequency in favorable and unfavorable environment will indicate that the only intrinsic growth rates are affected by the environment.

Here, we neglect the density dependent effects of the selection process by considering only frequency-dependent growth of different phenotypes. An extension of the model will be incorporation of the behavior of total population size as well, which may lead to richer evolutionary outcomes due to the dependence on absolute values of growth rates, thus, introducing a new degree of freedom. We also neglect the demographic noise in the model. The interplay between these two noises can lead to better understanding of the evolutionary process in nature, and provide new insights for combating the diseases.

## METHODS

The simulations of (4) are performed following [65] for piecewise-deterministic Markov processes. Namely, the next switching time is found by introducing an auxiliary quantity *g*(*t*) that satisfies 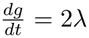, with *g*(0) = 0 initial conditions. Solving *g*(*t*_next_) = *−* ln *u* defines the next switching time, *t*_next_, where *u* is a random number sampled from uniform distribution *U* (0, 1). Once *t*_next_ is found, *g*(*t*) *→* 0. The next state of the environment is found by their respective stationary probabilities 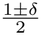 . For each tuples of model parameters, 10^4^ independent runs are carried out, with the initial conditions *x*(0) *∈ U* (0, 1), and 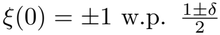 . For given values of model parameters, in each run *x*(*t*) is sampled at *t*_f_ = 100*τ* , where 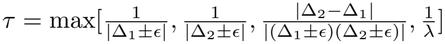 , where each component shows the timescale of convergence of each steady state of (4) (see SI section II.A for further details). The simulations are done both using Runge-Kutta numerical method with order of 4, precision taken at least min 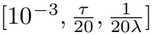 using double-precision floating point format and the Wolfram Mathematica platform [66].

For *Ψ*(*x*) in Fig.3B *N* = 5*×* 10^5^ simulations are carried on for each initial point *x*_0_ *∈* [*x*_+_ +*l, x*_*−*_ *− l*], with *l* = 10^*−*4^. Then it is divided by the number of overall simulations made for that particular setup. The standard deviation is then calculated using the formula 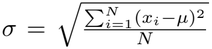 , where *N* is the total number of simulations for a given setup and *µ* is the mean hitting probability obtained from simulations. Wilson score [67] is then computed using 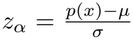 , where *p*(*x*) is the true binomial proportion of the outcomes for given *x*, and *α* = 0.05 corresponding to 95% confidence.

## COMPETING INTERESTS

The authors declare no competing interests.

## DATA AVAILABILITY

All study data are included in the article and/or SI Appendix. The code written in the course of this work was deposited through github https://github.com/BMEP-group/The-fate-of-social-dilemmas-under-randomly-fluctuating-growth-rates.git

## AUTHOR CONTRIBUTIONS

S.G.B. and K.H.C. conceptualized and designed the study. H.G.S, A.S.B and S.G.B. developed and performed the analytical and computational work. All authors analyzed and interpreted the data. S.G.B. and K.H.C. wrote the manuscript, with contributions from all authors. All authors have read and approved the final version of the manuscript.

## ACKNOWLEDGMENTS

This work was supported by the Higher Education and Science Committee of the Republic of Armenia under Research Program No. 24IRF/2-1C001. Kang Hao Cheong acknowledges support from the SPMS Collaborative Research Award 2025.

## Notes

### Competing Interest Statement

The authors have declared no competing interest.

