## Supplementary Information for "The fate of social dilemmas under randomly fluctuating growth rates"

### I. MAPPING LOTKA-VOLTERRA SYSTEM TO REPLICATOR DYNAMICS

In the model described by Lotka-Volterra for $L$ types, the dynamics of the population evolve according to the following system of differential equations:

$$\dot{n}_i = n_i \left( r_i + \sum_{j=1}^{L} \frac{n_j}{\sum_k n_k} a_{ij} \right), \text{ where } i \in 1, ..., L$$

Set the following substitutions: $x_i = \frac{n_i}{\sum_k n_k}$, $N = \sum_j n_j$ and $f_i = \sum_j (r_i + a_{ij}) x_j$. Then,

$$\begin{aligned}
\dot{x}_i &= \frac{d}{dt}\left(\frac{n_i}{N}\right) = \frac{\dot{n}_i N - n_i \dot{N}}{N^2} = \\
&= \frac{n_i}{N}\left( r_i + \sum_{j=1}^{L} \frac{n_j}{\sum_{k=1}^{L} n_k} a_{ij} \right) - x_i \sum_{j=1}^{L} \frac{\dot{n}_j}{N} = \\
&= x_i \left( \sum_{j=1}^{L} x_j (r_i + a_{ij}) - \sum_{j=1}^{L} x_j \sum_{k=1}^{L} x_k (r_j + a_{jk}) \right) = \\
&= x_i (f_i - \sum_{j=1}^{L} f_j x_j).
\end{aligned}$$

This is the ordinary differential equation for replicator dynamics. Thus for a population composition $\mathbf{x} \in \mathbb{R}^L, \forall k : x_k \geq 0$ and $\sum_k x_k = 1$ the fitness is defined as follows:

$$f_i(\mathbf{x}) = (\mathbf{A}\mathbf{x})_i = \sum_{j=1}^{L} (a_{ij} + r_i) x_j, \tag{1}$$

where $a_{ij} + r_i$ is the corresponding (shifted) entry in the game matrix $\mathbf{A}$, with $r_i$ being the intrinsic growth rate of type $i$.

In populations consisting of solely two traits ($L = 2$), following reduction is carried further (we focus on the frequency of the first trait, that is, $x \equiv x_1$):

$$\begin{aligned}
\dot{x} &= x(f_1 - x f_1 - (1 - x) f_2) = \\
&= x(1 - x)(f_1 - f_2).
\end{aligned}$$

Using the notation of the main paper following equation is obtained:

$$\dot{x} = x(1-x)(r_1 - r_2 + \Delta_2 + (\Delta_1 - \Delta_2)x), \tag{2}$$

whereby the parameters are defined as such: $\Delta_1 = a_{11} - a_{21}, \Delta_2 = a_{12} - a_{22}$.

#### I.A Analytic Solutions for the Deterministic Case

Note that the aforementioned differential equation (2) has an implicit closed-form solution. Rewrite as such ($\alpha = \Delta_2 + \epsilon, \beta = \Delta_1 - \Delta_2, \epsilon = r_1 - r_2$):

$$\begin{aligned} 1 &= \frac{dx}{dt} \cdot \frac{1}{x(1-x)(\alpha + \beta x)} = \\ &= \frac{dx}{dt}\left(\frac{1}{\alpha x} + \frac{1}{(\alpha+\beta)(1-x)} - \frac{\beta^2}{\alpha(\alpha+\beta)(\alpha+\beta x)}\right) = \\ &= \frac{d}{dt}\left(\frac{1}{\alpha}\log x - \frac{1}{\alpha+\beta}\log(1-x) - \frac{\beta}{\alpha(\alpha+\beta)}\log(\alpha+\beta x)\right). \end{aligned}$$

Subsequently, the following solution is obtained:

$$t + C = \frac{1}{\Delta_2 + \epsilon}\log x - \frac{1}{\Delta_1 + \epsilon}\log(1-x) - \frac{\Delta_1 - \Delta_2}{(\Delta_2+\epsilon)(\Delta_1+\epsilon)}\log|\Delta_2 + \epsilon + (\Delta_1 - \Delta_2)x|, \tag{3}$$

where $C \in \mathbb{R}$ is an integration constant.

### II. REPLICATOR DYNAMICS UNDER MARKOVIAN DICHOTOMOUS NOISE

Our model uses Markovian Dichotomous Noise, also known as telegraph noise. The noise element $\xi(t)$, which will be denoted below as $I_t$ to follow the convention of the book, takes discrete values (we denote $-$ and $+$ for unfavorable and favorable environments, respectively), transitions of which are following a Poisson process with fixed rates. In case of unequal rates, i.e. non-zero bias $\delta$, the transitions of corresponding states are calculated with different probabilities. Consequently, the noise $I_t$ takes values in $\{+\epsilon, -\epsilon\}$ and is a time-homogeneous Markov process with transition probabilities $P_{i,j}(t) = \text{Prob}(I_t = i \,|\, I_0 = j)$.

*Transition probabilities*

Let $\alpha \equiv \lambda(1+\delta)$ and $\beta \equiv \lambda(1-\delta)$ be the rates $+ \to -$ and $- \to +$ and set $\Lambda = \alpha + \beta$ (the relaxation rate; $\Lambda = 2\lambda$ in the parametrization above). The master equation reads

$$\frac{d}{dt}\begin{pmatrix} P_{-,j} \\ P_{+,j} \end{pmatrix} = \begin{pmatrix} -\beta & \alpha \\ \beta & -\alpha \end{pmatrix}\begin{pmatrix} P_{-,j} \\ P_{+,j} \end{pmatrix}. \tag{4}$$

The eigenvalues are 0 and $-\Lambda$ with eigenvectors $(\alpha, \beta)^T$ and $(1, -1)^T$, so

$$\begin{pmatrix} P_{-,j}(t) \\ P_{+,j}(t) \end{pmatrix} = c_1 \begin{pmatrix} \alpha \\ \beta \end{pmatrix} + c_2 e^{-\Lambda t}\begin{pmatrix} 1 \\ -1 \end{pmatrix}. \tag{5}$$

For $j = -$: $\alpha c_1 + c_2 = 1$, $\beta c_1 - c_2 = 0$, giving $c_1 = 1/\Lambda$, $c_2 = \beta/\Lambda$. For $j = +$: $\alpha c_1 + c_2 = 0$, $\beta c_1 - c_2 = 1$, giving $c_1 = 1/\Lambda$, $c_2 = -\alpha/\Lambda$. Collecting columns,

$$P(t) = \frac{1}{\Lambda}\begin{pmatrix} \alpha + \beta e^{-\Lambda t} & \alpha(1 - e^{-\Lambda t}) \\ \beta(1 - e^{-\Lambda t}) & \beta + \alpha e^{-\Lambda t} \end{pmatrix}. \tag{6}$$

As $t \to \infty$ both columns tend to the stationary distribution $P_s(+\epsilon) = \beta/\Lambda$, $P_s(-\epsilon) = \alpha/\Lambda$.

*Stationary distribution of the population composition*

The replicator equation with noise takes the form

$$\dot{X} = f(X) + I(t)\, g(X), \qquad f(x) = x(1-x)\big(\Delta_2 + (\Delta_1 - \Delta_2)x\big), \quad g(x) = x(1-x). \tag{7}$$

Let $P_\pm(x,t)$ be the probability density of $X$ jointly with $I_t = \pm\epsilon$. The forward Kolmogorov equations are

$$\partial_t P_+ = -\partial_x\big[(f + \epsilon g)P_+\big] - \alpha P_+ + \beta P_-,$$
$$\partial_t P_- = -\partial_x\big[(f - \epsilon g)P_-\big] + \alpha P_+ - \beta P_-.$$

In the stationary regime $\partial_t P_\pm = 0$. Adding the two equations shows that the total flux $(f + \epsilon g)P_+ + (f - \epsilon g)P_-$ is constant; zero-flux boundary conditions set it to zero, so

$$P_- = -\frac{f + \epsilon g}{f - \epsilon g} P_+.$$

(Positivity of both densities requires $f+\epsilon g$ and $f-\epsilon g$ to have opposite signs, so the stationary density is supported only on the interval between the fixed points of the two environments.) Substituting into the first stationary equation,

$$\frac{d}{dx}\big[(f+\epsilon g)P_+\big] = -\Big(\alpha + \beta\,\frac{f+\epsilon g}{f-\epsilon g}\Big)P_+,$$

so

$$P_+(x) = \frac{C}{|f+\epsilon g|}\exp\left[-\int^x \Big(\frac{\alpha}{f+\epsilon g} + \frac{\beta}{f-\epsilon g}\Big)du\right].$$

The marginal density is $\Pi_{\rm st} = P_+ + P_- = P_+\,\dfrac{-2\epsilon g}{f-\epsilon g}$, that is,

$$\Pi_{\rm st}(x) = \frac{1}{Z}\,\frac{g(x)}{\big|f(x)^2-\epsilon^2 g(x)^2\big|}\exp\left[-\int^x \Big(\frac{\alpha}{f+\epsilon g} + \frac{\beta}{f-\epsilon g}\Big)du\right]. \tag{8}$$

Evaluating the integral with $f\pm\epsilon g = x(1-x)\big(\Delta_2 \pm \epsilon + (\Delta_1-\Delta_2)x\big)$, $\alpha=\lambda(1-\delta)$, $\beta=\lambda(1+\delta)$, one obtains on the interval between the two fixed points

$$\begin{aligned}\Pi_{\rm st}(x) &= \frac{1}{Z}\,x^{-\frac{2\lambda(\Delta_2+\delta\epsilon)}{\Delta_2^2-\epsilon^2}-1}(1-x)^{\frac{2\lambda(\Delta_1+\delta\epsilon)}{\Delta_1^2-\epsilon^2}-1} \\ &\quad\times\big|(\Delta_1-\Delta_2)x+\Delta_2+\epsilon\big|^{\frac{\lambda(1-\delta)(\Delta_1-\Delta_2)}{(\Delta_1+\epsilon)(\Delta_2+\epsilon)}-1}\big|(\Delta_1-\Delta_2)x+\Delta_2-\epsilon\big|^{\frac{\lambda(1+\delta)(\Delta_1-\Delta_2)}{(\Delta_1-\epsilon)(\Delta_2-\epsilon)}-1}.\end{aligned} \tag{9}$$

Each of the four factors arises from one of the four simple zeros of $f^2-\epsilon^2 g^2$; the exponent is $-1$ (from the prefactor $g/|f^2-\epsilon^2 g^2|$) plus $\mp\alpha/c$ or $\mp\beta/c$ with $c$ the slope of the corresponding linear factor.

*Links to invasion rates of each phenotype*

In the main text we defined invasion rates $\Lambda_C = -(\Delta_1+\delta\epsilon)$ and $\Lambda_D = \Delta_2+\delta\epsilon$. Apparently those rates are also being used in the PDF (9), particularly in the exponents of $x$ and $1-x$. Specifically, (9) can be rewritten as such:

$$\begin{aligned}\Pi_{\rm st}(x) &= \frac{1}{Z}\,x^{-\frac{2\lambda\Lambda_D}{\Delta_2^2-\epsilon^2}-1}(1-x)^{-\frac{2\lambda\Lambda_C}{\Delta_1^2-\epsilon^2}-1} \\ &\quad\times\big|(\Delta_1-\Delta_2)x+\Delta_2+\epsilon\big|^{\frac{\lambda(1-\delta)(\Delta_1-\Delta_2)}{(\Delta_1+\epsilon)(\Delta_2+\epsilon)}-1}\big|(\Delta_1-\Delta_2)x+\Delta_2-\epsilon\big|^{\frac{\lambda(1+\delta)(\Delta_1-\Delta_2)}{(\Delta_1-\epsilon)(\Delta_2-\epsilon)}-1}.\end{aligned}$$

It is obvious that the signs of $\Lambda_C$ and $\Lambda_D$ deduce whether the integral of PDF is divergent near $x=0$ and $x=1$, as seen in the following analysis of the games.

### III. GAMES

In the main text three types of games are discussed. These are games based on a payoff matrix of the following form:

$$\mathbf{A} = \begin{bmatrix} P & T \\ S & R \end{bmatrix},$$

where the entry in $i$-th row and $j$-th column of the matrix is equal to the expected payoff received by a player employing strategy $i$ against strategy $j$. The names of the entries in the matrix are chosen according to the model of Prisoner's Dilemma merely for convenience. In evolutionary game theory instead of exact payoffs the entries indicate average payoffs, or average contributions to the fitness of the trait. One may use the fitness formula (1) obtained from the entries in the abovementioned matrix, thus yielding the following equality:

$$\mathbf{A} = \begin{bmatrix} P & T \\ S & R \end{bmatrix} \equiv \begin{bmatrix} a_{11} + r_1 & a_{12} + r_1 \\ a_{21} + r_2 & a_{22} + r_2 \end{bmatrix}.$$

Subsequently, in all of the games described below the reproductive advantage is essentially encoded in the values of expected payoffs.

#### III.A Hawk-Dove or Anti-Coordination Game

In the coexistence game, the carriers of the first trait are called **hawks**, while the carriers of the latter trait are labeled **doves** (due to their respective natures, hawks are willing to fight while doves yield and prefer peace). The parameters are chosen so that $T > R > S > P$, that is, the cost of fight is higher than the value of resource.

The essence of the game is as follows: a player chooses a strategy - whether to fight over a resource or retreat.

- Fighting yields payoff $\mathbf{P}$ if the opponent fights back (in a typical game of Hawk-Doves, $\mathbf{P} < 0$). If the opponent retreats, the player receives all the payoff for the resource, that is $\mathbf{T}$.

- The player retreats and receives payoff $\mathbf{R}$ if the opponent also chooses peace. However, if the opponent decides to fight back, the player will instantly give up and receive $\mathbf{S}$ (usually $\mathbf{S} = 0$).

Consider what happens when a resident hawk population is invaded by a small $0 < \omega \ll 1$ proportion of doves. Then their respective fitnesses are as follows:

$$\begin{aligned} f_1 &= P \cdot (1-\omega) + T \cdot \omega, \\ f_2 &= S \cdot (1-\omega) + R \cdot \omega, \\ f_1 - f_2 &= (1-\omega)(P-S) + \omega(T-R) = \\ &= (P-S) + \omega(T-R-P+S). \end{aligned}$$

Note that $P - S < 0$, therefore we may find $\omega^\star > 0$ such that $\forall \omega < \omega^\star$:

$$(P-S) + \omega(T-R-P+S) = f_1 - f_2 < 0,$$

which is equivalent to $\dot{x} < 0$, that is, the frequency of hawks will decrease. Hence, a full population of hawks is susceptible to invasion by an arbitrarily small number of doves.

In the contrary case, consider a population consisting mostly of doves. Then the fitnesses are such:

$$\begin{aligned} f_1 &= P \cdot \omega + T \cdot (1-\omega), \\ f_2 &= S \cdot \omega + R \cdot (1-\omega), \\ f_1 - f_2 &= \omega(P-S) + (1-\omega)(T-R) = \\ &= (T-R) + \omega(P-S-T+R), \end{aligned}$$

By the same logic, since $T - R > 0$, we may find another $\omega^\star > 0$ such that $\forall \omega < \omega^\star$:

$$f_1 - f_2 > 0,$$

that is, the proportion of hawks will increase.

Finding the stationary points of the original differential equation (2), it is easily seen that the aforementioned discussion points to the unique interior stationary state, which is also the evolutionary stable state of the game:

$$\bar{x} = \frac{r_1 - r_2 + \Delta_2}{\Delta_2 - \Delta_1}. \tag{10}$$

*Analysis of the System*

For the analysis of the stochastic system further conditions must be held by the parameters in order for the system to consist of switching between two games of coexistence. Note that the value of the parameter $\epsilon \equiv r_1 - r_2$ (that is, the differential of intrinsic growth rates) offsets the stable point $\bar{x}$ by an amount of $\frac{\epsilon}{\Delta_2 - \Delta_1}$. Thus, for the novel fixed points caused by the fluctuations to be within the feasible region $(0, 1)$ the following inequality is required:

$$\epsilon < \min(-\Delta_1, \Delta_2),$$

which represents an upper bound on the amplitude of noise. Then we denote stationary points as such:

$$x_\pm = \frac{\Delta_2 \pm \epsilon}{\Delta_2 - \Delta_1}.$$

Note that both of these points are the only interior fixed points in their respective environments, specifically, they are the evolutionary stable states of their corresponding games.

*Behaviour at the endpoints*

Take $\delta = 0$. The only possible singularities are at $x_\pm$, with exponents $\frac{\lambda(\Delta_1 - \Delta_2)}{(\Delta_1 \pm \epsilon)(\Delta_2 \pm \epsilon)} - 1$. Because $\Delta_1 < 0$ and $\epsilon < -\Delta_1$, we have $(\Delta_1 - \Delta_2)/\big((\Delta_1 \pm \epsilon)(\Delta_2 \pm \epsilon)\big) > 0$, so both exponents exceed $-1$ and $\Pi_{\text{st}}$ is always integrable. The density vanishes at both endpoints iff both exponents are positive, i.e.

$$\lambda > \max\left[\frac{(\epsilon - \Delta_1)(\Delta_2 - \epsilon)}{\Delta_2 - \Delta_1}, \ \frac{(\Delta_1 + \epsilon)(\Delta_2 + \epsilon)}{\Delta_1 - \Delta_2}\right]. \tag{11}$$

This agrees with the intuition that at large switching rates the system behaves like the time-averaged deterministic system and does not reach the endpoints. If $|\Delta_1| \neq |\Delta_2|$, $\lambda$ can be chosen so that the density vanishes at only one endpoint.

*Extrema and the M-shape distribution*

Setting $\partial_x \ln \Pi_{\text{st}} = 0$ in (8) (with $\alpha = \beta = \lambda$) and multiplying through by $(f^2 - \epsilon^2 g^2)/(2\lambda)$ gives

$$f + \frac{1}{\lambda}\Big(ff' - \tfrac{1}{2}\frac{g'}{g}f^2 - \tfrac{1}{2}\epsilon^2 g g'\Big) = 0, \tag{12}$$

with $f = x(1-x)h(x)$, $h(x) = \Delta_2 + (\Delta_1 - \Delta_2)x$, $g = x(1-x)$. Explicitly,

$$x(1-x)h - \frac{\epsilon^2}{2\lambda}x(1-x)(1-2x) + \frac{1}{\lambda}x(1-x)h\Big[(1-x)h - xh + x(1-x)(\Delta_1 - \Delta_2)\Big]$$
$$- \frac{1}{2\lambda}x(1-x)(1-2x)h^2 = 0.$$

An M-shape density has zeros at both endpoints, so (11) must hold, together with three roots of (12) in $(x_-, x_+)$ (two maxima separated by a minimum). The region in parameter space where this occurs is found numerically and shown in the region plot of the main text.

#### III.B Prisoner's Dilemma or Game of Dominance

In the game of dominance, the following inequality holds for the initial parameters of the game matrix: $T > R > P > S$. The traits are named **defectors** and **cooperators**, respectively. The essence of the game is as follows:

A player is facing a dilemma - whether to cooperate or defect.

- Cooperation yields payoff **R** (reward for cooperation) if the opponent chooses the same, and gives **S** (sucker's payoff) otherwise, that is, the opponent cheats or defects.

- If the player defects instead, they receive payoff **T** (temptation to cheat) if the opponent chooses to cooperate, or they receive **P** (punishment) if the opponent decides to defect alike.

According to the setup of the parameters, mutual cooperation is better than mutual defection. However, for each individual player defection is the best option, hence the dilemma occurs. In game theory, mutual defection is the Nash equilibrium, while in evolutionary game theory, mutual defection is the evolutionary stable strategy, and dominance of defectors ($x^\star = 1$) is the unique evolutionary stable state, because $\forall x : f_1 > f_2$.

*Equilibria and Stability Analysis*

Following the convention above, we note the following stationary points:

$$x = 0, \qquad x = 1, \qquad x_\pm = \frac{\Delta_2 \pm \epsilon}{\Delta_2 - \Delta_1}.$$

Further below is discussed all possible cases for the initial parameters of the game ($\Delta_1$, $\Delta_2$, $\epsilon$).

- Case $\Delta_1 > \Delta_2$:

| Case | Regime | Env. | $x=0$ | $x=1$ | $x_\pm$ |
|---|---|---|---|---|---|
| 1 | $\epsilon > \Delta_1 > \Delta_2$ | $+\epsilon$ | $U$ | $S$ | $x_+ < 0$ $(S)$ |
| | | $-\epsilon$ | $S$ | $U$ | $x_- > 1$ $(S)$ |
| 2 | $\Delta_1 > \epsilon > \Delta_2$ | $+\epsilon$ | $U$ | $S$ | $x_+ < 0$ $(S)$ |
| | | $-\epsilon$ | $S$ | $S$ | $x_- \in (0,1)$ $(U)$ |
| 3 | $\Delta_1 > \Delta_2 > \epsilon$ | $+\epsilon$ | $U$ | $S$ | $x_+ < 0$ $(S)$ |
| | | $-\epsilon$ | $U$ | $S$ | $x_- < 0$ $(S)$ |

- Case $\Delta_2 > \Delta_1$:

| Case | Regime | Env. | $x=0$ | $x=1$ | $x_\pm$ |
|---|---|---|---|---|---|
| 4 | $\epsilon > \Delta_2 > \Delta_1$ | $+\epsilon$ | $U$ | $S$ | $x_+ > 1$ $(U)$ |
| | | $-\epsilon$ | $S$ | $U$ | $x_- < 0$ $(U)$ |
| 5 | $\Delta_2 > \epsilon > \Delta_1$ | $+\epsilon$ | $U$ | $S$ | $x_+ > 1$ $(U)$ |
| | | $-\epsilon$ | $U$ | $U$ | $x_- \in (0,1)$ $(S)$ |
| 6 | $\Delta_2 > \Delta_1 > \epsilon$ | $+\epsilon$ | $U$ | $S$ | $x_+ > 1$ $(U)$ |
| | | $-\epsilon$ | $U$ | $S$ | $x_- > 1$ $(U)$ |

Here $S$ denotes stable and $U$ denotes unstable. Equilibria outside $[0, 1]$ are mathematically valid roots but are not physically admissible states.

The interior equilibrium $x_-$ belongs to $(0, 1)$ if and only if

$$\min(\Delta_1, \Delta_2) < \epsilon < \max(\Delta_1, \Delta_2).$$

The equilibrium $x_+$ never belongs to $(0, 1)$.

For first case, the exponent of the term $(1 - x)$ in stationary PDF becomes less than $-1$ due to the ordering $\epsilon > \Delta_1$, thus inducing a peak therein at the fixed point 1. The same reasoning applies to case 5 (which is discussed in the main text) and case 4. For case 2 the reasoning fails due to an unstable interior point induced in the unfavorable environment.

Case 3 and 6 fail to be of any interest, because the only admissible stable point is the dominance of defectors.

*Mean First Passage Time*

In the symmetric case of Prisoner's Dilemma, particularly zero bias and symmetric environmental fluctuations, the analysis of the stationary PDF fails to convey any peculiarities in the behavior of the game. Nonetheless, another interesting aspect worth considering is the mean first passage time (MFPT). We are interested in MFPT to boundary $x_0 = 1 - \eta$ (for a sufficiently small $\eta$), taken so because the trajectory never actually reaches $x = 1$. Hence for a specified boundary $x_0$ let $T_\xi(x_0)$ define the mean first passage time when the particle hits $x_0$:

$$
\begin{aligned}
T_\xi(x_0) = (1 - \lambda\Delta t) T_\xi\left(x_0 + [f(x_0) + g(x_0)\xi]\Delta t\right) \\
+ \lambda \Delta t\, T_{-\xi}(x_0) + \Delta t + o(\Delta t).
\end{aligned} \tag{13}
$$

After expanding the Eq. (45) using Taylor series and neglecting the terms of order $o(\Delta t^2)$ we obtain two differential equations for

$$
f(x) = x(1-x)\big(\Delta_2 + (\Delta_1 - \Delta_2)x\big), \quad g(x) = x(1-x).
$$

$$
\begin{aligned}
[f(x_0) + g(x_0)]\frac{dT_+}{dx} - \lambda(T_+ - T_-) = -1, \\
[f(x_0) - g(x_0)]\frac{dT_-}{dx} + \lambda(T_+ - T_-) = -1.
\end{aligned} \tag{14}
$$

Following the same steps for derivation of hitting probability we get:

$$
\begin{aligned}
F(x_0) = A \exp\left[-2\lambda \int^{x_0} \frac{f(u)}{g(u)^2 - f(u)^2}\, du\right] + \exp\left[-2\lambda \int^{x_0} \frac{f(u)}{g(u)^2 - f(u)^2}\, du\right] \times \\
\times \left[\int^{x_0} \exp\left[2\lambda \int^{z} \frac{f(u)}{g(u)^2 - f(u)^2} du\right] * \frac{2g(z)}{f(z)^2 - g(z)^2}\, dz\right],
\end{aligned} \tag{15}
$$

where $F(x_0) = T_+(x_0) - T_-(x_0)$. Again, define

$$
H(x_0) = \exp\left[-2\lambda \int_{x_+}^{x_0} \frac{f(u)}{g(u)^2 - f(u)^2}\, du\right]. \tag{16}
$$

Then $F(x_0) = AH(x_0) + H(x_0)\int^{x_0} \frac{2g(z)}{(f(z)^2 - g(z)^2)H(z)} dz$.

From the Eqs. (29) and (14) the differential equations for hitting time in plus and minus environments will be:

$$
\begin{aligned}
\frac{dT_+}{dx} &= \frac{\lambda F(x_0) - 1}{g(x_0) + f(x_0)}, \\
\frac{dT_-}{dx} &= \frac{\lambda F(x_0) + 1}{g(x_0) - f(x_0)}.
\end{aligned}
$$

Integrating from $x_-$ to $x_0$, we obtain

$$
\begin{aligned}
T_+(x_0) = \lambda \int_{x_-}^{x_0} \frac{F(y) - 1}{g(y) + f(y)}\, dy + B_1, \\
T_-(x_0) = \lambda \int_{x_-}^{x_0} \frac{F(y) + 1}{g(y) - f(y)}\, dy + B_2,
\end{aligned} \tag{17}
$$

where $B_1$ and $B_2$ are integration constants. The boundary conditions will be:

$$T_+(x_-) - T_-(x_-) = -\frac{1}{\lambda},$$
$$T_+(1-\eta) = 0.$$

The meaning is that it takes $\frac{1}{\lambda}$ time to switch from + to - environment on average, and at point $1-\eta$ the hitting time is 0, as we already hit the desired point.

The average hitting time for $x_0$ will be:

$$T(x_0) = \frac{1}{2}\left[T_+(x_0) + T_-(x_0)\right].$$

For the deterministic case, the following expression can be found from the original solution (3) (which may be calculated for $+\epsilon, -\epsilon$ and $\epsilon = 0$):

$$t_\epsilon(x_0) = C + \frac{1}{\Delta_2+\epsilon}\ln(x_0) - \frac{1}{\Delta_1+\epsilon}\ln(1-x_0) - \frac{\Delta_1-\Delta_2}{(\Delta_1+\epsilon)(\Delta_2+\epsilon)}\ln|\Delta_2+\epsilon+(\Delta_1-\Delta_2)x_0|. \tag{18}$$

Hence set the mean hitting time for the deterministic case as such:

$$\tau_\epsilon(x_0) = t_\epsilon(1-\eta) - t_\epsilon(x_0).$$

Now fix $x_0 = \frac{x_-+1}{2}$ and compute MFPT to $x = 1-\eta$ for different switching rates, $\Phi(\lambda) = T_\lambda(x_0)$. For $\lambda \approx 0$ the initial environment is chosen at first with probability 1/2 (in symmetric case; for biased case it is $\frac{1-\delta}{2}$ and $\frac{1+\delta}{2}$) and never changes, so roughly half of the trajectories never reach $1-\eta$ in the bad environment. Hence $\tau_{-\epsilon}(x_0) = +\infty$ and the average over the initial state is likewise:

$$\tau(x_0) = \tfrac{1}{2}\tau_\epsilon(x_0) + \tfrac{1}{2}\tau_{-\epsilon}(x_0) = +\infty.$$

Note that for sufficiently large $\lambda$ the environment averages out, so $\Phi(\lambda) \to \tau_0(x_0)$. Moreover, it follows from the reasoning above that $\Phi(\lambda) \to +\infty$ as $\lambda \to 0^+$. Consequently, if there exists some $\lambda_1 > 0$ such that $\Phi(\lambda_1) < \tau_0(x_0)$, continuity of $\Phi(\lambda)$ implies the existence of a local minima $\lambda^* \in (0, +\infty)$.

Two modes for the function $\Phi(\lambda)$ therefore occur, as shown in Fig. 1B: in the first mode, the function decreases monotonically in $\lambda$ and stays strictly above $\tau_0(x_0)$. In the second mode, the function decreases to a local minimum at some $\lambda^*$ and then increases and approaches the deterministic time $\tau_0(x_0)$ while staying strictly below it. Our numerical experiments show that the parameter $\Delta_1$ contributes the most to the switching of the modes, namely for a sufficiently small $\Delta_1$ the function $\Phi(\lambda)$ takes the first form and then transitions to the latter form as the value

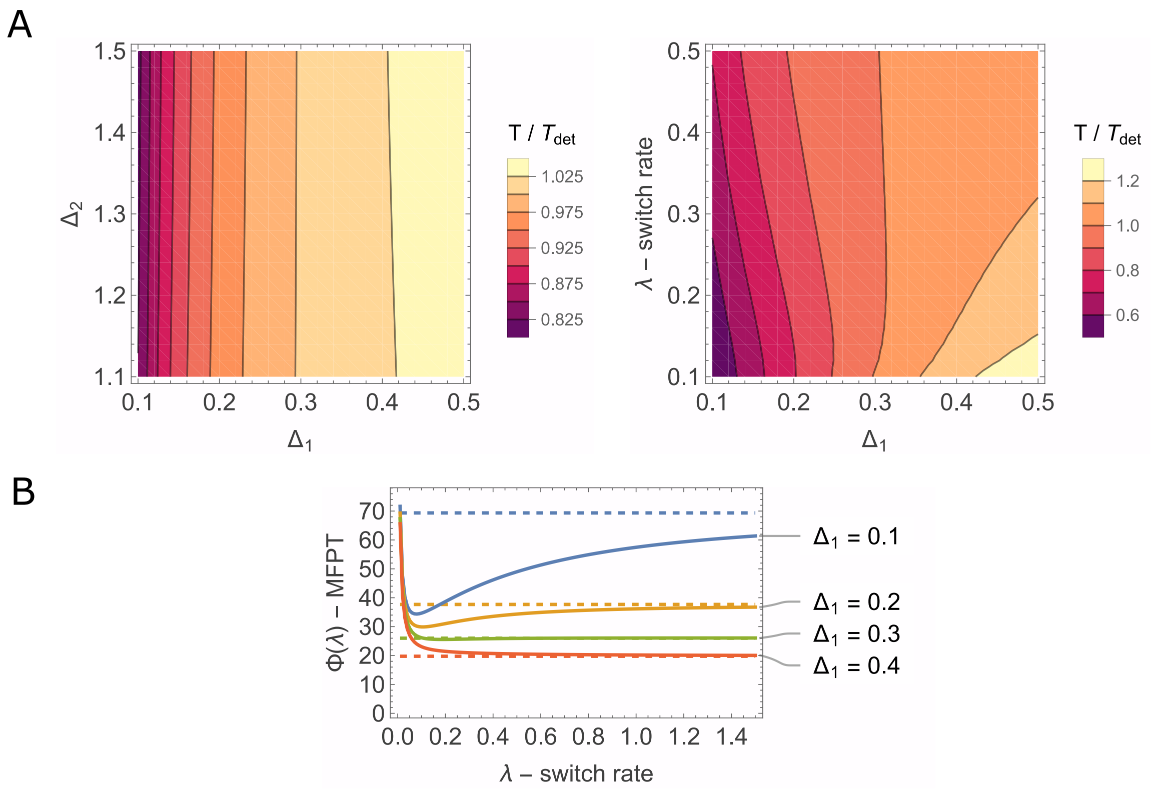


FIG. 1: **Numerical MFPT for the Prisoner's Dilemma game.**

**A)** Heatmaps of the ratio of the MFPT in the fluctuating environment to the deterministic hitting time $\tau_0$ (the averaged environment, $\epsilon = 0$). In the left heatmap, $\Delta_1$ affects the ratio more strongly than $\Delta_2$. For some parameter values the ratio is smaller than 1, i.e. the MFPT in the fluctuating environment is shorter than the deterministic hitting time. The remaining parameters are $\lambda = 1.0$, $\epsilon = 0.75$ for the left heatmap and $\Delta_2 = 1.0$, $\epsilon = 0.75$ for the right heatmap.

**B)** Dependence of the MFPT $\Phi(\lambda) = T_\lambda(x_0)$ on the switching rate $\lambda$ for several values of $\Delta_1$ ($x_0 = 1 - \eta$, $\eta = 10^{-4}$, other parameters: $\epsilon = 0.75$, $\Delta_2 = 1.0$). Solid lines show the MFPT; dashed lines show the deterministic hitting time for the corresponding parameters. The shape of $\Phi(\lambda)$ changes at a threshold near $\Delta_1 = 0.3$ (green line): below it, $\Phi$ decreases monotonically with $\lambda$; above it, $\Phi$ is non-monotonic, decreasing to a minimum at $\lambda^*$ and then increasing towards the deterministic value.

of $\Delta_1$ increases (see Fig. 1B). In order to find the relation between parameters and the mode of the function $\Phi(\lambda)$, numerical solutions may be used to determine the regions of parameters where $\Phi(\lambda)$ behaves correspondingly.

*Biased Environment*

In the main text we examine the regime $\Delta_2 > \epsilon > \Delta_1$ and obtain the requirements for $\delta$ that change the outcome of the game. The case $\epsilon > \Delta_2 > \Delta_1$ is quite similar to the one mentioned above with the only difference that under unfavorable environment instead of $x = x_-$ the stable fixed point is $x = 0$. In order to find the divergence and convergence conditions for the case $\epsilon > \Delta_2 > \Delta_1$ we need to examine the exponents of the terms $x$ and $1-x$, namely $\frac{-2\lambda(\Delta_2+\delta\epsilon)}{\Delta_2^2-\epsilon^2} - 1$ and $\frac{2\lambda(\Delta_1+\delta\epsilon)}{\Delta_1^2-\epsilon^2} - 1$. Solving the inequalities

$$\frac{-2\lambda(\Delta_2+\delta\epsilon)}{\Delta_2^2-\epsilon^2} - 1 > -1,$$
$$\frac{2\lambda(\Delta_1+\delta\epsilon)}{\Delta_1^2-\epsilon^2} - 1 > -1,$$

for $\delta$, we find that $\delta < -\frac{\Delta_1}{\epsilon}$ so around $1-x$ the PDF is integrable and $\delta > -\frac{\Delta_2}{\epsilon}$ for integration around $x$. As, $\Delta_2 > \Delta_1$, $\delta$ must be less than $-\frac{\Delta_2}{\epsilon}$ so that in both ends the stationary PDF is integrable. In order to get bell shaped distribution we solve these inequalities:

$$\frac{-2\lambda(\Delta_2+\delta\epsilon)}{\Delta_2^2-\epsilon^2} - 1 > 0,$$
$$\frac{2\lambda(\Delta_1+\delta\epsilon)}{\Delta_1^2-\epsilon^2} - 1 > 0,$$

which yields $\frac{-2\lambda\Delta_2-\Delta_2^2+\epsilon^2}{2\lambda\epsilon} < \delta < \frac{\Delta_1^2-\epsilon^2-2\lambda\Delta_1}{2\lambda\epsilon}$. Moreover, these can be rewritten as such:

$$\lambda > \max\left[\frac{\epsilon^2-\Delta_2^2}{2(\Delta_2+\delta\epsilon)}, \frac{\epsilon^2-\Delta_1^2}{-2(\Delta_1+\delta\epsilon)}\right]. \tag{19}$$

This corresponds with the intuition that higher switching rates average out the environment, resulting in a bell-shape distribution. Failing to satisfy either of the conditions results in a peak at the corresponding boundary.

*Unequal advantages in intrinsic growth rates*

The purpose of asymmetric noise is to give full advantage to C and observe whether the situation changes, at all. Firstly, we have to define the stationary PDF for the asymmetric case, where $\Delta = \Delta_-$ and $\Delta_+ = 0$. Now the Fokker-Plank equation becomes:

$$0 = -\frac{d}{dx}\left[fP_+\right] - \alpha P_+ + \beta P_-,$$
$$0 = -\frac{d}{dx}\left[(f+\Delta_- g)P_-\right] + \alpha P_+ - \beta P_-.$$

Using the same steps as in II.B we find that the stationary PDF for asymmetric case is:

$$\begin{aligned}\Pi_{\mathrm{st}-}(x) = \quad & \frac{1}{Z_1} x^{-\frac{2\lambda\Delta_2-\lambda\epsilon}{\Delta_2(\Delta_2-\epsilon)}-1} \times (1-x)^{\frac{2\lambda\Delta_1-\lambda\epsilon}{\Delta_1(\Delta_1-\epsilon)}-1} \times \\ & \times |(\Delta_1-\Delta_2)x+\Delta_2|^{\frac{\lambda(\Delta_1-\Delta_2)}{\Delta_1\Delta_2}-1} \times |(\Delta_1-\Delta_2)x+\Delta_2-\epsilon|^{\frac{\lambda(\Delta_1-\Delta_2)}{(\Delta_1-\epsilon)(\Delta_2-\epsilon)}-1}.\end{aligned} \tag{20}$$

In order to make the stationary PDF integrable around $x = 1$ following condition must hold:

$$\frac{2\lambda\Delta_1-\lambda\epsilon}{\Delta_1(\Delta_1-\epsilon)} - 1 > -1, \tag{21}$$

which yields to the inequality below:

$$\epsilon > 2\Delta_1. \tag{22}$$

Note that we may do the following substitutions:

$$\begin{aligned}\Delta_i' &= \Delta_i - \frac{\epsilon}{2}, \text{ for } i \in \{1,2\}, \\ \epsilon' &= \frac{\epsilon}{2}.\end{aligned}$$

Analyzing the signs of this new system, we find that $\Delta_1' < 0 < \epsilon' < \Delta_2'$ if the condition (22) is satisfied, hence the observed case is the Hawk-Dove game. Therefore the condition (22) ensures that the game is mapped to Hawk-Dove case with symmetric noise.

Furthermore, in the asymmetric noise case we can observe M-shape distribution, as well. Then the PDF must vanish (hence be integrable) at both endpoints $x_- = \frac{\Delta_2-\epsilon}{\Delta_2-\Delta_1}$ and $x = 1$:

$$\frac{\lambda(\Delta_1-\Delta_2)}{(\Delta_1-\epsilon)(\Delta_2-\epsilon)} - 1 > 0, \tag{23}$$

$$\frac{2\lambda\Delta_1-\lambda\epsilon}{\Delta_1(\Delta_1-\epsilon)} - 1 > 0. \tag{24}$$

If both conditions are satisfied, the shape of the distribution is either an M-shape or a bell shape. If either of the conditions fail, a peak is observed there. The condition (24) is especially dangerous because it can create a non-integrable divergence, thus invalidating the conclusion that this is a Hawk-Dove game. To get M-shape distribution, one should look at the equation of the extrema for $\Pi_{\mathrm{st}-}(x)$:

$$\begin{aligned}&\Delta_2^2 + 2\Delta_2\lambda - \Delta_2\epsilon - 4x^3(\Delta_1-\Delta_2)^2 + 3x^2(\Delta_1-\Delta_2)(\Delta_1-3\Delta_2+\epsilon)+ \\ &+ 2\Delta_1 x(2\Delta_2+\lambda-\epsilon) - 2\Delta_2 x(3\Delta_2+\lambda-2\epsilon) - \lambda\epsilon = 0,\end{aligned} \tag{25}$$

which must have three distinct roots. Numerical solutions provide the exact regions for parameters to encounter M-shape distribution, shown in Fig.2D of the main text.

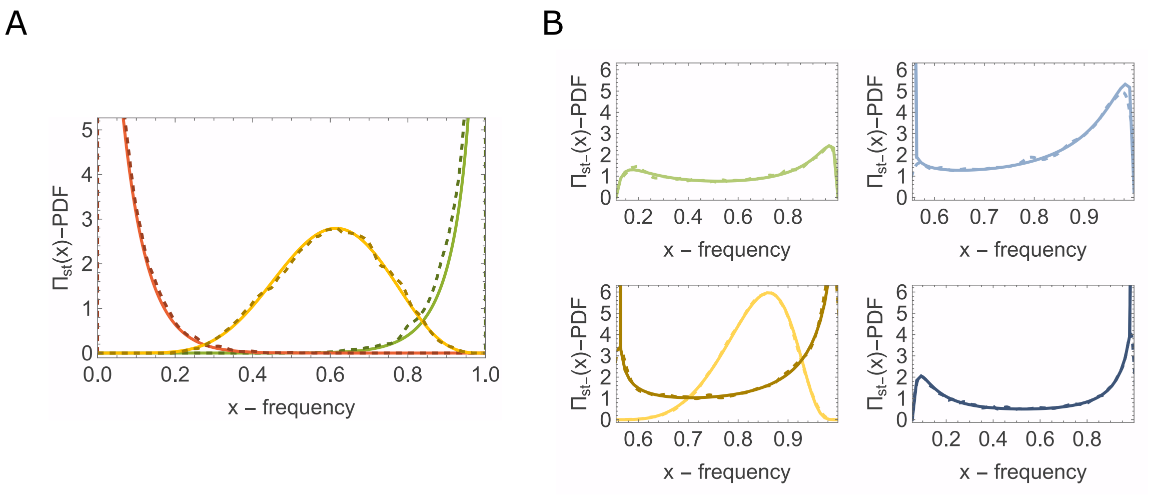


FIG. 2: **Stationary PDFs of the Prisoner's Dilemma game for case 4 with non-zero bias ($\epsilon > \Delta_2 > \Delta_1$) and for asymmetric noise.**

**A)** Case 4, $\epsilon > \Delta_2 > \Delta_1$. At zero bias ($\delta = 0$) the PDF is concentrated at a single boundary point and no non-trivial distribution exists, whereas a non-zero bias yields several distinct shapes: a bell shape (yellow; $\delta = -0.3$) and a PDF vanishing at one of the boundary points (red; $\delta = -0.48$, and green; $\delta = -0.18$). The remaining parameters are $\Delta_1 = 0.1$, $\Delta_2 = 0.3$, $\epsilon = 0.6$, and $\lambda = 10.0$.

**B)** Asymmetric noise (unequal advantages). Solid lines show the PDF given by the analytic solution (20); dashed lines show the distribution reconstructed from numerical simulations. The colors of the plots correspond to the colored regions of the heatmap in Fig. 3D of the main text. As in the Hawk-Dove game, an M-shape distribution is obtained for particular parameter values (top-left, green; $\epsilon = 0.9$, $\lambda = 0.17$). The PDF may also vanish at one of the boundary points (top-right, light blue; $\epsilon = 0.5$, $\lambda = 0.17$, and bottom-right, dark blue; $\epsilon = 0.95$, $\lambda = 0.1$), and bell-shaped and U-shaped distributions are possible (bottom-left, light yellow, bell-shaped; $\epsilon = 0.5$, $\lambda = 1.0$, and dark yellow, U-shaped; $\epsilon = 0.5$, $\lambda = 0.1$). The remaining parameters are $\Delta_1 = 0.1$ and $\Delta_2 = 1.0$.

#### III.C Stag-Hunt or Coordination Game

In the game of stag-hunt, two players decide whether to coordinate or act alone, both given the options to hunt for either a stag (adult male deer) or a hare, hence the traits are accordingly labeled as **stag** and **hare**. The parameter ordering is as follows: $P > S > R > T$. The interactions are defined as follows:

- If both players cooperate and hunt for a stag, their payoff will be the highest, **P**. However, if the first player hunts a stag alone, while the other chooses to hunt a hare, the first player will most likely fail, thus they receive the lowest payoff **T**.

- If the player decides to hunt for a hare alone, they receive **S**. If both players coordinate and hunt for a hare together, they will be rewarded **R**.

The game has two pure Nash equilibria. In the evolution of population dynamics, the system has two stable stationary points which correspond to the said equilibria.

Moreover, the original equation (2) has an interior unstable fixed point, which is the following:

$$\bar{x} = \frac{r_1 - r_2 + \Delta_2}{\Delta_2 - \Delta_1} \tag{26}$$

*Analysis of the System*

Similar to Hawk-Dove, we find that the system exhibits two equilibria, which are merely shifted versions of (26), by $\frac{\epsilon}{\Delta_2 - \Delta_1}$. Condition $\epsilon < min(\Delta_1, -\Delta_2)$ ensures that both of the points belong to the interval $(0, 1)$. Hence, we find the following unstable fixed points corresponding to respective environments:

$$x_\pm = \frac{\Delta_2 \pm \epsilon}{\Delta_2 - \Delta_1},$$

where, unlike the Hawk-Dove case, $x_+ < x_-$.

Consider an initial point $x(0) = x_0$. Based on the value of this initial state, we may categorize the evolution of the trajectory:

- $x_0 < x_+$: due to the influence of two unstable fixed points to the right of $x_0$, further trajectory of the point will just approach the nearest stable point, that is 0. Subsequently, all the trajectories for which $x(0) < x_+$ will converge to 0.

- $x_0 > x_-$: same reasoning applies here; all the trajectories satisfying $x(0) > x_-$ will converge to 1.

- For $x_0 \in (x_+, x_-)$ the trajectory will lie within the same interval until it passes either boundary, after which it will just follow the behavior described above.

*Derivation of the Hitting Probability*

In the Stag-Hunt game, all trajectories approach either leftmost $x = 0$ or rightmost $x = 1$ boundary, meaning that the analysis of stationary PDF is not useful. Therefore, we resort to analyzing hitting probabilities dependent on the initial position of the particle.

Our process evolves according to

$$\dot{x}(t) = f(x(t)) + g(x(t))\xi(t), \tag{27}$$

where

$$f(x) = x(1-x)\big(\Delta_2 + (\Delta_1 - \Delta_2)x\big), \quad g(x) = x(1-x).$$

Let $\psi_\xi(x)$ denote the probability that a particle starting at position $x$ with orientation $\xi$ reaches the right boundary $x = x_-$ before reaching the left boundary $x = x_+$.

Consider the evolution over a small time interval $\Delta t$. With probability $1 - \lambda\Delta t$, the orientation does not flip, and the particle moves to

$$x + [f(x) + g(x)\xi(t)]\,\Delta t.$$

With probability $\lambda\Delta t$, the orientation flips from $\xi$ to $-\xi$. Therefore,

$$\begin{aligned}\psi_\xi(x) = (1 - \lambda\Delta t)\psi_\xi\left(x + [f(x) + g(x)\xi]\Delta t\right)\\ + \lambda\Delta t\,\psi_{-\xi}(x) + o(\Delta t).\end{aligned} \tag{28}$$

Expanding the first term on the right-hand side to first order gives

$$\begin{aligned}\psi_\xi(x) = (1 - \lambda\Delta t)\left[\psi_\xi(x) + [f(x) + g(x)\xi(t)]\frac{d\psi_\xi}{dx}\Delta t\right]\\ + \lambda\Delta t\,\psi_{-\xi}(x) + o(\Delta t).\end{aligned}$$

Neglecting terms of order $(\Delta t)^2$ and higher, and cancelling $\psi_\xi(x)$ from both sides, we obtain

$$[f(x) + g(x)\xi]\frac{d\psi_\xi}{dx} + \lambda[\psi_{-\xi}(x) - \psi_\xi(x)] = 0. \tag{29}$$

For the two orientations, this yields the coupled equations

$$[f(x) + g(x)]\frac{d\psi_+}{dx} - \lambda(\psi_+ - \psi_-) = 0, \tag{30}$$

$$[f(x) - g(x)]\frac{d\psi_-}{dx} + \lambda(\psi_+ - \psi_-) = 0. \tag{31}$$

The boundary conditions follow from the direction of motion at the boundaries. A particle at $x = x_-$ with orientation $+1$ immediately exits through the right boundary, while a particle at $x = x_+$ with orientation $-1$ immediately exits through the left boundary. Thus,

$$\psi_+(x_-) = 1, \qquad \psi_-(x_+) = 0. \tag{32}$$

*Transformation to sum and difference variables*

To solve the coupled equations, define

$$Q(x) = \psi_+(x) + \psi_-(x), \qquad F(x) = \psi_+(x) - \psi_-(x). \tag{33}$$

It follows that

$$\psi_+ = \frac{Q+F}{2}, \qquad \psi_- = \frac{Q-F}{2}.$$

Adding Eqs. (30) and (31), the switching terms cancel, giving

$$f(x)\frac{dQ}{dx} + g(x)\frac{dF}{dx} = 0. \tag{34}$$

Subtracting Eq. (31) from Eq. (30) gives

$$g(x)\frac{dQ}{dx} + f(x)\frac{dF}{dx} = 2\lambda F(x). \tag{35}$$

From Eq. (34),

$$\frac{dQ}{dx} = -\frac{g(x)}{f(x)}\frac{dF}{dx},$$

where this intermediate expression is understood away from points where $f(x) = 0$. Substituting directly into Eq. (35) gives

$$\left[f(x) - \frac{g(x)^2}{f(x)}\right]\frac{dF}{dx} = 2\lambda F(x).$$

Multiplying by $f(x)$ and rearranging yields the first-order equation

$$[f(x)^2 - g(x)^2]\frac{dF}{dx} - 2\lambda f(x)F(x) = 0. \tag{36}$$

*Solution for the difference function*

Rearranging Eq. (36), we find

$$\frac{1}{F(x)}\frac{dF}{dx} = \frac{2\lambda f(x)}{f(x)^2 - g(x)^2} = -\frac{2\lambda f(x)}{g(x)^2 - f(x)^2}.$$

Integrating from $x_+$ to $x$ gives

$$\ln\left|\frac{F(x)}{F(x_+)}\right| = -2\lambda \int_{x_+}^{x} \frac{f(u)}{g(u)^2 - f(u)^2}\,du.$$

Consequently, the solution can be written as

$$F(x) = A\exp\left[-2\lambda \int_{x_+}^{x} \frac{f(u)}{g(u)^2 - f(u)^2}\,du\right], \tag{37}$$

where $A$ is an integration constant.

For convenience, define

$$H(x) = \exp\left[-2\lambda \int_{x_+}^{x} \frac{f(u)}{g(u)^2 - f(u)^2}\, du\right]. \tag{38}$$

Then $F(x) = AH(x)$.

*Recovery of $\psi_+$ and $\psi_-$*

From Eqs. (34) and (35),

$$\frac{dQ}{dx} = -\frac{g(x)}{f(x)}\frac{dF}{dx}.$$

Using Eq. (36), an equivalent expression that avoids division by $f(x)$ is

$$\frac{dQ}{dx} = \frac{g(x)f(x)}{g(x)^2 - f(x)^2}\, 2\lambda \frac{F(x)}{f(x)} = \frac{2\lambda g(x)F(x)}{g(x)^2 - f(x)^2}.$$

Alternatively, differentiating the individual functions directly in Eqs. (30) and (31), and using $\psi_+ - \psi_- = F$, gives

$$\frac{d\psi_+}{dx} = \frac{\lambda F(x)}{g(x) + f(x)},$$
$$\frac{d\psi_-}{dx} = \frac{\lambda F(x)}{g(x) - f(x)}.$$

Integrating from $x_+$ to $x$, we obtain

$$\psi_+(x) = \lambda \int_{x_+}^{x} \frac{F(y)}{g(y) + f(y)}\, dy + B_1, \tag{39}$$
$$\psi_-(x) = \lambda \int_{x_+}^{x} \frac{F(y)}{g(y) - f(y)}\, dy + B_2, \tag{40}$$

where $B_1$ and $B_2$ are integration constants.

At $x = x_+$, the definition of $F$ and the boundary condition $\psi_-(x_+) = 0$ imply

$$F(x_+) = \psi_+(x_+) - \psi_-(x_+) = \psi_+(x_+).$$

Since $H(x_+) = 1$, we have $A = \psi_+(x_+) = B_1$. Therefore,

$$B_1 = A.$$

Using $F = \psi_+ - \psi_-$ at $x = x_+$ also gives

$$B_1 - B_2 = A,$$

and hence $B_2 = 0$.

It follows that

$$\psi_+(x) = A\left[1 + \lambda \int_{x_+}^{x} \frac{H(y)}{g(y)+f(y)}\,dy\right], \tag{41}$$

$$\psi_-(x) = A\lambda \int_{x_+}^{x} \frac{H(y)}{g(y)-f(y)}\,dy. \tag{42}$$

Finally, applying the remaining boundary condition $\psi_+(x_-) = 1$ to Eq. (41) determines $A$:

$$A = \left[1 + \lambda \int_{x_+}^{x_-} \frac{H(y)}{g(y)+f(y)}\,dy\right]^{-1}. \tag{43}$$

Thus, the hitting probabilities are

$$\psi_+(x) = \frac{1 + \lambda \displaystyle\int_{x_+}^{x} \frac{H(y)}{g(y)+f(y)}\,dy}{1 + \lambda \displaystyle\int_{x_+}^{x_-} \frac{H(y)}{g(y)+f(y)}\,dy}, \tag{44}$$

$$\psi_-(x) = \frac{\lambda \displaystyle\int_{x_+}^{x} \frac{H(y)}{g(y)-f(y)}\,dy}{1 + \lambda \displaystyle\int_{x_+}^{x_-} \frac{H(y)}{g(y)+f(y)}\,dy}, \tag{45}$$

where $H(y)$ is defined in Eq. (38).

If the initial orientation is equally likely to be $+1$ or $-1$, the probability of exiting through the right boundary is

$$\psi(x) = \frac{1}{2}\left[\psi_+(x) + \psi_-(x)\right]. \tag{46}$$

In the Fig.3 of the main text, the hitting probabilities are evaluated numerically.
